# Alphafold 3 guided insights into the Importin β - Importin 7 heterodimer interaction and its binding to Histone H1

**DOI:** 10.1101/2025.08.04.668392

**Authors:** Piotr Neumann, Olexandr Dybkov, Henning Urlaub, Ralf Ficner, Achim Dickmanns

## Abstract

The nuclear import of H1 linker histones is facilitated by a heterodimer of the transport receptors Importinβ (Impβ) and Importin7 (Imp7). While both importins can individually interact with H1, only their preassembled hetero-dimer enables its proper binding for translocation through the nuclear pore. The interaction between Imp7 and Impβ is mediated by a short stretch of residues in the C-terminal region of Imp7, which plays a key role in Imp7 activation by Impß. This interaction is allosterically regulated by Impβ, finely tuning the activity of the Impβ/Imp7 heterodimer. A complete model of the Impβ:Imp7:H1 complex was predicted by Alphafold3 (AF3) and subsequently validated using cross-linking (X-link) data, isothermal titration calorimetry (ITC), and pull-down experiments, providing robust support for the model’s accuracy. This model positions the globular domain of H1 within the central cavity of Imp7, in agreement with cross-linking data. Refinement of this atomic model against a previously published cryo-EM map demonstrated significantly improved correspondence compared to the earlier interpretation, which placed the H1 globular domain within Impβ. This enhanced structural consistency further substantiates the accuracy of the AI-driven prediction. Detailed analysis identified the nucleoporin-like binding (NlB) region of Imp7 as a short stretch interacting with the outer surface of Impβ. This interaction mode implies that regulation of FG-binding site accessibility on Impβ, mediated by the FXFG nucleoporin motives within the Imp7 NlB region, likely modulates transport pathways and/or enhances the efficiency of H1 nuclear import.

## INTRODUCTION

A prerequisite for functional compartmentalization of eukaryotic cells is the existence of transport systems that facilitate the exchange of molecules between distinct compartments. Among them, the nucleus is enclosed by a double layered membrane, which is punctuated by aqueous channels formed by nuclear pore complexes (NPCs). These large supramolecular structures are built up by 30 different nuclear pore proteins, so called nucleoporins (Nups), that occur in multiples of eight and form various stable subcomplexes ultimately resulting in the structural assembly of an eightfold symmetric ring-like or cylindrical overall structure ^1–6^. Among these, 10 Nups contribute to the formation of a selective permeability barrier within the central aqueous channel of the NPC. This barrier, the mechanism of which is explained by various models^7^, is created by intrinsically disordered regions (IDRs) of those Nups. These IDRs include the FG repeat regions, named after the Phe-Gly residues that are spaced by linkers typically comprising 12-20 residues ^8–11^. The resulting meshwork allows smaller solutes, ions and small molecules to freely diffuse between the compartments. In contrast larger proteins typically require the assistance of specific transport receptors. They bind the larger cargo and facilitate interactions with the FG repeats, enabling their passage through the permeability barrier.

Three main classes of transport receptors facilitate nuclear-cytoplasmic transport. The first class is involved in mRNA export, utilizing Mex67/Mtr2 TAP/p15 ^12–15^. The second class specializes in importing the small GTPase Ran named NTF2 ^16–18^. The third class comprises the karyopherin beta superfamily, responsible for shuttling cargo between the nucleus and the cytosol in both directions. Within this family, the karyopherins are divided into exportins and importins, although this distinction is not absolute, as some karyopherins function in both roles. Examples of the latter include Importin 13 ^19–21^, Exportin 4 (XPO4) ^22,23^ and Exportin 7 (XPO7)^24,25^.

All karyopherins share a structural framework composed of a tandem array of helix-turn-helix motifs, so called HEAT repeats (named after **H**untingtin, elongation factor 3 (**E**F3), protein phosphatase 2**A** (PP2A), yeast kinase **T**OR1). HEAT repeats (typically 19–24) are arranged in a stacked, superhelical structure with the A-helix forming the convex outer surface and the B-helix formed concave inner surface. The concave side mediates substrate recognition, often harboring binding sites. The first 7–8 repeats bind the small GTPase Ran ^26,27^, which acts as a sensor for the receptor-cargo complex location to ensure transport directionality. Cargo-binding sites, overlapping with the Ran-binding domain, recognize specific motifs such as nuclear localization signals (NLS, e.g., PY-motif) for import and nuclear export signals (NES, e.g., classic NES) for export^28,29^.

Nuclear import is not solely dependent on direct receptor-cargo interactions; two additional mechanisms expand cargo specificity. Importin-β (Impβ) broadens its cargo range via adaptor proteins like Importin-α and Snurportin 1, which bridge cargo binding through an N-terminal Importin-β-binding (IBB) domain. Receptor dimerization further diversifies cargo transport, e.g. CRM1 homo-dimers facilitate HIV RNA export ^30,31^, while heterodimers, such as Impβ-Importin7 (Imp7), transport nuclear factors (HIV integrase, histone H1, L4, L6)^32–36^. These cargos are characterized by small size, intrinsically disordered regions, and basic residue enrichment, reflecting their role as DNA/RNA-binding proteins and necessitating acidic receptor patches for charge compensation. Remarkably, Imp7 is predicted to contain a long acidic loop within its 19–20 HEAT repeats^37^, and exhibits a more complex structural arrangement than Impβ due to additional insertions within and between repeats.

Histones are essential for DNA packaging into nucleosomes and higher-order chromatin, particularly during S-phase when demand surges. The nucleosome core comprises an octamer of H2A, H2B, H3, and H4, wrapping ∼146 bp of DNA ^38^. Histone H1, a linker histone, secures DNA entry and exit sites on the nucleosome ^39^. In contrast to linker histones, the core histones (H2A, H2B, H3, H4) are 100–140 residues long, sharing a conserved histone fold (∼70 residues) and lysine-rich N-terminal tails (20–40 residues) ^40^. H2A and H2B also possess short C-terminal tails of 12 or 13 residues. Nuclear import of these histones is mediated by Importin-β superfamily members: Importin 9 (Imp9) transports H2A/H2B, while Importin 4 (Imp4) imports H3/H4 ^41–46^. While individual transport receptors suffice for dimeric or monomeric core histones, the import of the linker histone H1 is more complex. H1, typically exceeds 200 aa in length and features a distinct structure with an extended, lysine-rich C-terminal tail (80–100 residues) and a central globular winged-helix domain (the core domain consisting of three α-helices followed by an antiparallel pair of β-strands) ^47–50^. Its high positive charge differentiates it from core histones.

Nuclear import of H1 is achieved by the hetero-dimeric complex of Impβ:Imp7. Biochemical and structural investigations have identified the regions involved in this interaction (Supplementary Fig. S1) and the steps of complex assembly. Imp7 binds to Impβ via its C-terminal disordered region, which contains FG motifs and interacts with HEAT repeats 5-8 of Impβ ^34–37^. This domain, referred to by various names such as Importinβ binding site or IBB7^35,36^, will hereafter be termed the NlB (Nucleoporin-like Binding) region, as this term most aptly describes its function. The interaction of the NlB region of Imp7 with Impβ is a prerequisite for the formation of an import competent ternary complex with H1 ^34,36^. The binding regions of H1 have been mapped to the central part of Impβ and the C-terminal half of Imp7. The previously published incomplete cryo-EM structure of the Imp7:Impβ:H1 complex positions the globular core domain of H1 (residues 22-95) within the concave inner curvature of Impβ, suggesting that the remaining regions of the histone interact primarily with Imp7 ^37^. This placement is based on *in vitro* import studies demonstrating that the core globular domain of H1 by itself is imported by Impβ ^35^, which aligns with the observation that RanGTP triggers the release of Impβ from it. Conversely, pull-down experiments indicate that Imp7 exhibits a higher binding affinity for H1 compared to Impβ and that both H1 and an importin β-binding (IBB) domain can simultaneously associate with the Imp7-Impβ complex^34^. This evident discrepancy is further highlighted by the aforementioned structural studies, which reported cross-linking data that contradicted the proposed arrangement of the Imp7:Impβ:H1 complex (PDB ID: 6N88) but were consistent with our own cross-linking results of a similar complex. Recent advances in AI-driven structure prediction tools inspired us to revisit the ternary import complex using AlphaFold3 (AF3)^51,52^ and to reassess the arrangement of Imp7, Impβ, and H1 and their interaction sites. In contrast to the previously deposited cryo-EM structure (PDB 6N88, ^37^), the revised arrangement is strongly supported by cross-link data from two complexes that differ only in their histone components, the published Imp7:Impβ:H1.0 (human histone, ^37^) and Impβ:Imp7: H1.11L (chicken histone, this study) (Fig. 1). By integrating the available cryo-EM map (EMD-0366 ^37^) with the AF3-predicted model, a revised structural arrangement of the ternary complex Impβ:Imp7:H1.0 was determined and validated. This revised model, (Fig. 1B, C), reveals swapped positions of the Imp7 and Impβ proteins, leading to its re-designation as Impβ:Imp7:H1.0. A detailed analysis of the interaction between the C-terminal NlB region of Imp7 and Impβ was conducted, and the results corroborated by ITC measurements of individual point mutations, providing additional evidence for the accuracy of the reassessed Impβ:Imp7:H1.0 complex arrangement.

**Fig. 1.**
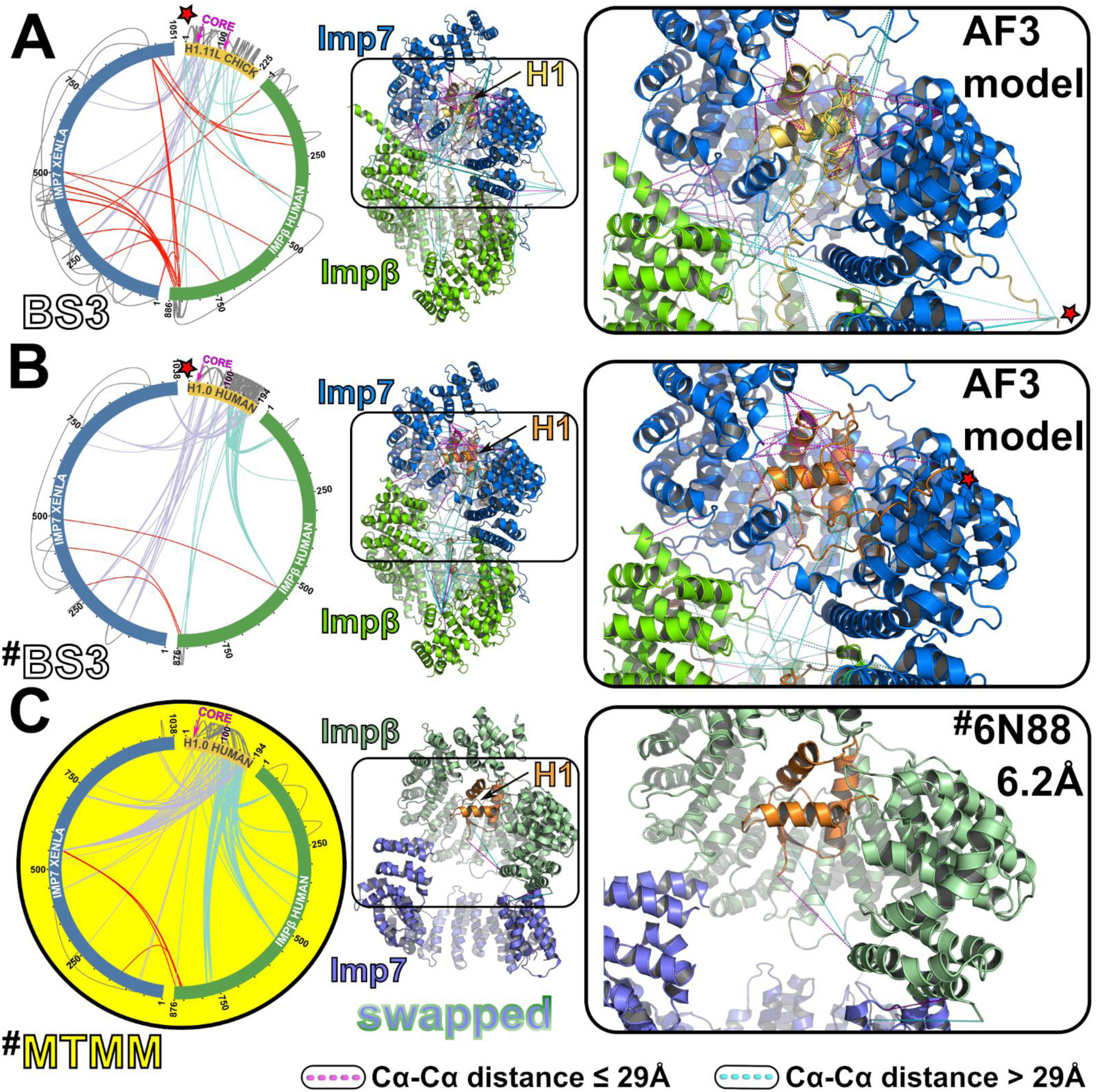
Cross-linking experiments confirm the structural models generated by AlphaFold 3 (AF3). Imp7 is depicted in blue, Impβ in green and H1 in yellow-orange. **A.** Impβ:Imp7:H1.11L-chick BS3 data, **B**. and **C.** Impβ:Imp7:H1.0-hs BS3 and MTMM data, respectively, taken from Ivic et al 2019. Left panels: Proteins indicated as bars in a circular plot created with xiVIEW, with cross-links shown in red for Imp7-Impβ, Violet for Imp7-H1 and Cyan for H1-Impβ. Middle panels: Overall models of the ternary complex of Impβ:Imp7:H1 (chicken, AF3)(top), Impβ:Imp7:H1 (human, AF3)(middle) and Imp7:Impβ:H1 (human, PDB id: 6N88)(bottom). Right panel: enlarged views of the boxed regions indicated in the middle panels. Cross-linking distances are represented as dotted lines: red for distances ≤ 29 Å and cyan for distances > 29 Å.

## RESULTS

The Impβ:Imp7 heterodimer serves as the exclusive nuclear import receptor for rpL4, rpL6, and linker histone H1 ^34–36^. The binding sites for Imp7 and H1.0 on Impβ have been mapped to HEAT repeats 4–14 and 6–9, respectively (Supplementary Fig. S1A) ^33–36^. These overlapping regions form the core interaction domain of Impβ. Imp7 associates with its cargo upon binding to Impβ, and its C-terminal region (aa 665–1001) is predicted to contain a unique structural feature namely an extended acidic loop near its end ^36^. Remarkably, the Impβ:Imp7 interaction critically depends on a ∼30-residue C-terminal segment of Imp7^35^. The energetics of these interactions have been characterized via isothermal titration calorimetry (ITC)^36^ and pull-down experiments revealed that key residues within the NlB region, particularly the GGxxF and FxFG motifs, are essential for binding ^37^. These motifs likely interact within clefts formed by the outer A-helices of HEAT repeats 5-6-7 of Impβ ^37^. However, structural insights of these interactions remained incomplete, as the cryo-EM model of the Imp7:Impβ:Histone-H1.0 complex (PDB ID: 6N88) lacks critical molecular details due to missing side chains and large unresolved segments of Imp7 (residues 1–282, 371–380, 871–1038). To address this, we focused on the NlB region of Imp7 (residues 1002-1038), identified based on secondary structure predictions ^36^ to elucidate its interaction with Impβ at the atomic level. The predicted interaction, modeled using AlphaFold3, was validated against experimental data.

### AlphaFold3 predicted Impβ:Imp7:H1 complex structures support experimental cross-link data

Atomic models of Impβ:Imp7:H1.11L-chick^AF3^ and Impβ:Imp7:H1.0-hs^AF3^ complexes, differing only in the sequences of histone molecules (chicken versus human), were predicted using the AlphaFold3 (AF3) server. In order to enhance structural diversity in predictions, 100 Impβ:Imp7:H1.11L-chick complexes were generated through 20 individual runs with differing random seeds. For the Impβ:Imp7:H1.0-hs complex, five predictions (1 run) were obtained. These 100 predictions (models) of Impβ:Imp7:H1.11L-chick^AF3^ complexes (Supplementary Fig. S7) displayed consistent assembly of Impβ and Imp7, with calculated RMSDs ranging from 0.607 Å (14,279 atoms) to 3.265 Å (13,240 atoms) when compared to a randomly selected reference prediction. An identical assembly of Impβ and Imp7 was observed in the predicted 5 Impβ:Imp7:H1.0-hs ^AF3^ complexes, with calculated RMSDs ranging from 1.405 Å (14,816 atoms) to 2.975 Å (15,030 atoms) relative to the same reference model. Remarkably, the predicted positions of the core globular domain of both histone molecules (H1.11L-chick: region 41–114; H1.0-hs: region 20–97), which were not considered during superposition, remained consistent, as reflected by the small RMSD values. For H1.11L-chick, RMSDs ranged from 1.016 Å (529 atoms) to 3.063 Å (529 atoms), while for H1.0-hs, RMSDs ranged from 3.287 Å (454 atoms) to 3.552 Å (454 atoms). All predicted AF3 models were subsequently evaluated for their compatibility with available cross-linking data: Impβ:Imp7:H1.11L-chick^AF3^ (this study, Supplementary Table S3) and Impβ:Imp7:H1.0-hs^AF3^ ^37^. The evaluation involved measuring the Cα-Cα distances between cross-linked residues, using a threshold of 29 Å for the BS3 crosslinker. The predicted 100 Impβ:Imp7:H1.11L-chick^AF3^ models demonstrated consistency with 21-33 (out of 49) intermolecular cross-links between H1.11L and the Impβ:Imp7 heterodimer, while the Impβ:Imp7:H1.0-hs^AF3^ models aligned with 15-30 cross-links (Fig. 1, Supplementary Tables S4 or S5). The cross-linking pattern supports an intertwined orientation of the Imp7 and Impβ superhelices, with their C-terminal halves closely associated (Supplementary Tables S3, S4, S5). The N-terminal HEAT repeats of Impβ, which harbor the conserved “CRIME” region essential for RanGTP binding, remain highly accessible. Comparison of cross-linking data of Impβ:Imp7:H1.11L-chick (this study) and Impβ:Imp7:H1.0-hs ^37^ revealed only marginal changes in the interaction pattern in the presence of two histone molecules (Fig. 1A, B). Given that chick H1.11L, like human H1, consists of nearly 30% lysines, more than half of the inter molecular cross-links were assigned to the flexible regions of H1 (residues 1–40 and 120–225), while the remaining cross-links involved the structurally conserved H1 core domain comprising only ∼ 70 residues. The H1 core domain interacts exclusively with Imp7, while the flexible regions primarily associate with Impβ (Fig. 1A, B). This suggests that the conserved WGH motif of H1 may play a key role in stabilizing ternary complex assembly and disassembly, a hypothesis explored further in subsequent sections. Unexpectedly, the cross-link based interaction pattern between the H1.0-hs core domain and Imp7, as predicted by AF3, is not supported by the deposited cryo-EM structure of the Imp7:Impβ:H1.0 complex (PDB ID: 6N88) (Fig. 1)^37^, except for a single proximity between residue 97 and Impβ residue 867. This discrepancy prompted a reassessment of the original cryo-EM model, particularly regarding the positioning of Impβ and Imp7, which appear to be swapped. Taken together, our analyses suggests that the initial structural model of Imp7:Impβ:H1 should be reconsidered, raising questions about the accuracy of its original interpretation.

### The Impβ:Imp7:H1.0-hs^AF^^3^ and Imp7:Histone-H1.0^AF^^3^ models reveal improved fit to the experimental cryo-EM maps

Two sets of complexes, including the deposited structures Imp7:Impβ:H1.0-hs (PDB ID: 6N88) and Impβ:H1.0-hs (PDB ID: 6N89), as well as AF3-predicted models with swapped positions of Impβ and Imp7 molecules (Impβ:Imp7:H1.0-hs^AF3^ and Imp7:H1.0-hs^AF3^), were subjected to re-refinement against their corresponding cryo-EM maps (EMD-0366 and EMD-0367, respectively). The thorough re-refinement process demonstrated that the AF3-derived atomic models ^37^, provide a substantially improved fit to the respective experimental cryo-EM maps. This evaluation utilized multiple global validation metrics, including the FSC curve (model versus map), FSCaverage, and model-map correlation coefficient (CC) (Fig. 2, Supplementary Fig. S5, Supplementary Table S6). These comprehensive validation indicators unequivocally confirm that the positions of Impβ and Imp7 in the deposited structures were swapped (Supplementary Fig. S8 and S9).

**Fig. 2.**
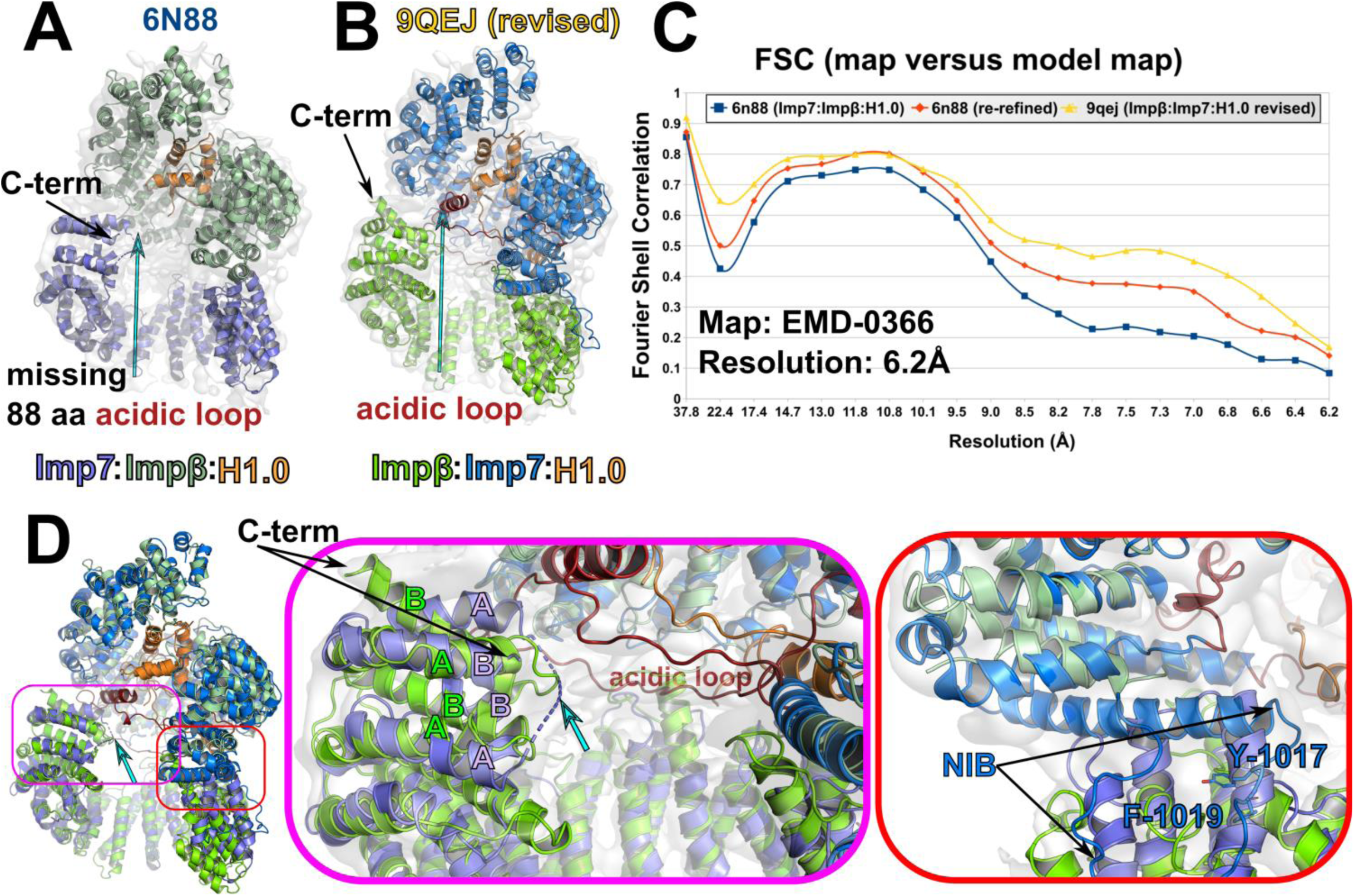
Structural refinement using the available cryo-EM map (EMD-0366) reveals significantly improved fit of the AlphaFold 3 (AF3)-generated model of the Impβ:Imp7:H1 (human). **A.** Structure of Imp7:Impβ:H1 (PDB id: 6N88). **B.** Refined and remodeled AF3-based structure of Impβ:Imp7:H1 reveals additional structural features present in the map (see D). **C.** The Fourier shell correlation (FSC) analysis comparing the refined atomic models and the cryo-EM map. **D.** Superposition of the revised structural model (Impβ:Imp7:H1) and the original interpretation (PDB id: 6N88). Magnifications, indicated by the boxes, are depicted on the right. Structures colored as in Figure 1.

The revised Impβ:Imp7:H1.0-hs structure shows substantial similarity to the Impβ:Imp7:H1.0-hs^AF3^ model, despite a high RMSD of 5.9 Å between 1.853 compared Cα atoms (Supplementary Fig. S8). This apparent discrepancy stems from the more compact conformations adopted by the Imp7 and Impβ molecules, encoded by the cryo-EM map. The Impβ:Imp7:H1.0-hs complex reveals a slightly reduced number of Lys residue pairs consistent with intra and inter molecular BS3 cross-linking data when compared with the AF3 model: 40 crosslinks versus 46 at a stringent 29 Å Cα-Cα distance cutoff or 46 crosslinks versus 48, when a more permissive 33 Å cutoff was applied (Table S4). The revised structural models have been deposited in the PDB under accession codes 9QEJ and 9QF0 for the Impβ:Imp7:Histone-H1.0 and Imp7:Histone-H1.0 complexes, respectively.

### The revised structure provides new insights into the Impβ:Imp7:H1.0-hs complex

In the Imp7:Impβ:H1.0-hs structure (PDB id: 6N88), the truncated model of Imp7, lacking residues 1–282, 371–380, 871–1038, was built into a cryo-EM map region corresponding to the significantly shorter Impβ molecule (Supplementary Fig. S9). This misassignment led to the erroneous conclusion that Imp7 adopts a conformation highly similar to Impβ, implying an atypical Impβ conformation within the heterodimeric transport receptor complex. However, structural comparison of the revised models of Impβ (residues 167–876) and full-length Imp7 (residues 1–1038) contradicts these findings. The refined Impβ model, excluding its flexible N-terminal 166 residues, adopts the characteristic curved solenoid architecture observed in multiple Impβ crystal structures (PDB IDs: 8GCN, 2Q5D, 1UKL). Strikingly, this conformation closely resembles the configuration of HEAT repeats 5–16 of the revised Imp7 model (Supplementary Fig. S9), while both the N-terminal and C-terminal HEAT repeat regions of Imp7 exhibit greater conformational flexibility and diverge substantially from Impβ. Moreover, the four N-terminal HEAT repeats of Impβ remain poorly resolved in the cryo-EM map, consistent with their intrinsic structural flexibility. The revised Impβ:Imp7:H1.0-hs model corrects previously unassigned or misinterpreted regions that were incompatible with the experimental cryo-EM map (Fig. 2), providing a more accurate structural representation of the ternary import complex. For instance, the last modeled HEAT repeat of Imp7, situated adjacent to the truncated acidic loop segment, was misoriented in the original model (6N88), with helices A and B incorrectly swapped (Fig. 2D middle). The correction aligns the Imp7 architecture with canonical HEAT repeat topology, further corroborating the revised structural arrangement. Additionally, reassessment of unassigned regions in the EMD-0366 map reveals detailed interactions involving the Imp7 NlB region (Fig. 2D right, Fig. 3A) and the H1.0-hs binding site on Imp7, which engages multiple HEAT repeats and the acidic loop – a feature absent in the original interpretation (Fig. 2B/D, 3B). The revised Impβ:Imp7:H1.0-hs assembly provides the first full-length structural model of Imp7, confirming its unique features within the importin family, including the aforementioned acidic loop and elongated HEAT repeat helices, which were previously predicted from sequence analysis but remained structurally unverified.

**Fig. 3.**
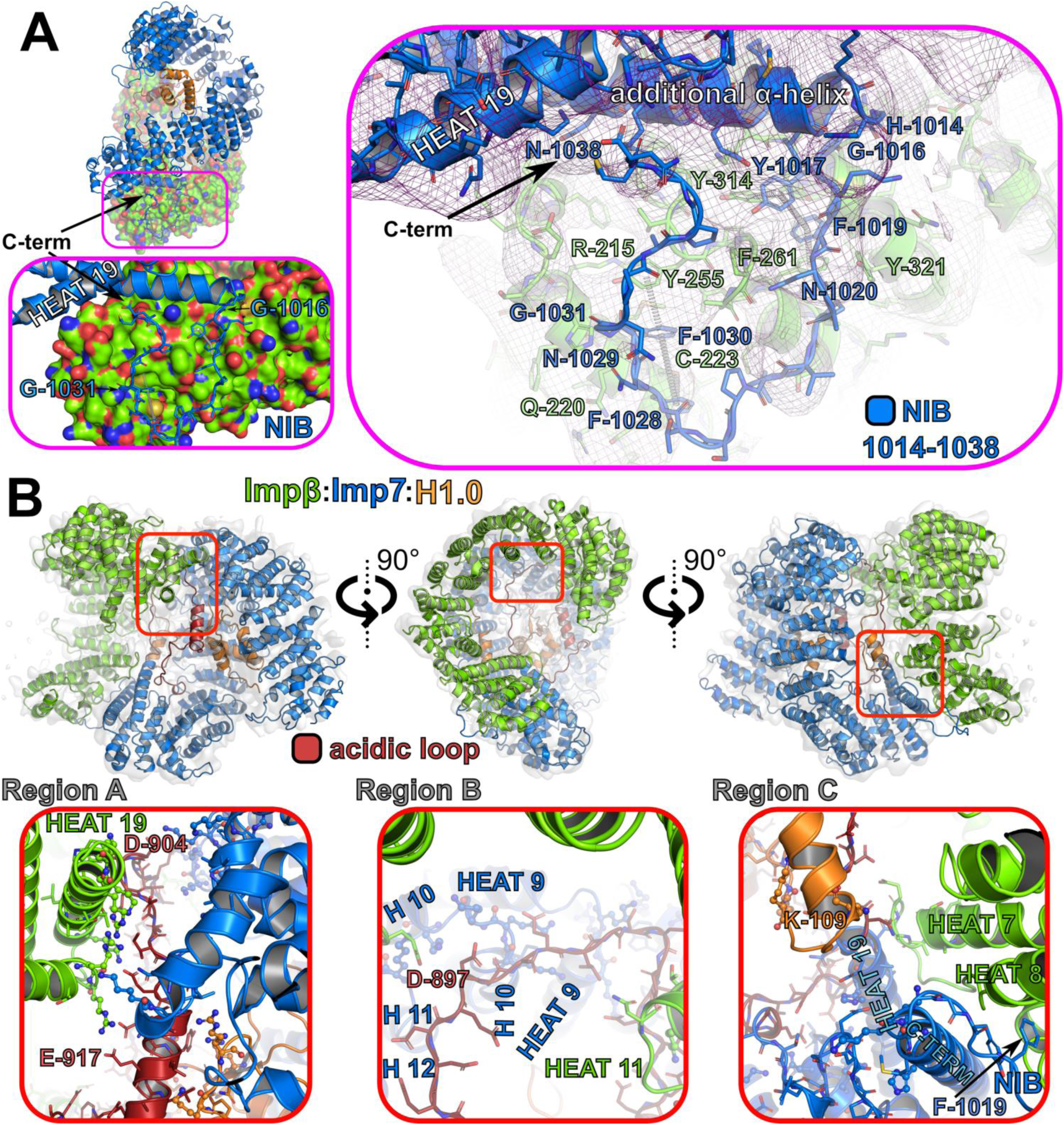
Structural features of Imp7 and Impβ interaction. **A.** Overall structure of the Impβ:Imp7:H1 complex, with a boxed region highlighting the NlB binding site on Impβ. The magnified view (below) details this interaction, alongside the corresponding cryo-EM map (right). The surface of Impβ is rendered and color-coded by atom type. Protein coloring follows the scheme in Figure 1. **B.** The acidic loop (red) in the C-terminal region of Imp7 acts as a structural barrier between H1 and Impβ. The top panels display three different views, while the lower panels present magnified views of the red-boxed regions above.

### Structure description

The integration of two cross-linking datasets (Impβ:Imp7:H1.11L-chick and Impβ:Imp7:H1.0-hs) along with the comprehensive validation of the latter complex against the cryo-EM map provides strong evidence that the H1 globular body is closely attached to the inner curvature of Imp7. Impβ consists of the well-characterized superhelical stacking of 19 HEAT repeats, where HEAT 19 helix A contains a notable kink, splitting it and giving it the appearance of an ARM-repeat 53. Its overall structure is solenoidal, consistent with established Impβ structures (Supplementary Fig. S6). The HEAT repeats are arranged in a uniform right-handed superhelical configuration, interrupted at specific points by extended loops of more than ten residues. Key deviations include: a helical insertion between HEAT 2 and 3 observed in crystal structures, an elongated loop spanning the gap between a long and a short helix of HEAT 7 and 8, and the acidic loop of 12 residues located between A and B helices of HEAT 8. The right-handed superhelical structure of Imp7, extending from HEAT1 to HEAT17, also exhibits the characteristic antiparallel orientation of the A and B helices, along with a well-defined hydrophobic interaction pattern within the helices, as described for canonical HEAT repeats ^53,54^. While most HEAT repeats are connected by short linkers between the two helices within each repeat, there are exceptions featuring longer insertions. The short acidic loop located in HEAT 6 of Imp7 (AL_H6_) comprises 19 residues, of which 9 are charged, though only two residues carry a positive charge. Similarly, HEAT 8 of Imp7 features a 31-residue insertion, with a charge distribution similar to standard interloop regions, consisting of around 25% charged residues. HEAT 19, on the other hand, comprises a highly extended 74-residue insertion, referred to as acidic loop (AL), containing 40 charged residues, though only one of these residues is positively charged (Fig. 3B). In comparison, the acidic loop of Impβ, located within HEAT 6, is negatively charged at 58% (7 of its 12 residues), illustrating clear differences in the length, charge composition and functional specificity between the two transport receptors. The last two HEAT repeats in Imp7 retain the defining characteristics of HEAT repeats, particularly the hydrophobic interaction between the A– and B-helices. However, they deviate from the standard tandem array observed in earlier repeats. Following HEAT 17, a 20-residue insertion begins with a helix, causing a roughly 90° shift in HEAT 18 relative to HEAT 17. HEAT 18 and HEAT 19 as well as an additional helix then form a distorted five-helix bundle, which extends into the NlB region (Fig. 3A). These distinctive structural features, including the longer insertions and deviations in the helical arrangements, highlight Imp7’s unique role within the β-karyopherin family. They underscore its specialized function in nuclear transport, influencing both its interactions with binding partners and its conformational flexibility.

### Heterodimeric assembly of Imp7 and Impβ

The two transport receptors, Impβ and Imp7, interact in a C-terminal-to-C-terminal annealed conformation, involving the last 13 HEAT repeats of both proteins. Their antiparallel superhelices intercalate to form an extended superhelical structure with a U-shaped central cavity, providing a shielded environment for cargo binding and chaperoning – a hallmark of their transport function. The HEAT repeats 1-4 of Impβ, absent in the revised cryo-EM structure, further extend the superhelical architecture (Fig. 1A). The N-terminal six HEAT repeats of both receptors, forming the conserved Ran-GTP binding CRIME domain, remain readily accessible^26,27^.

The Imp7-Impβ interaction is stabilized by four distinct regions. The first region, Region A, involves the long acidic loop (AL) in HEAT 19, which forms multiple salt bridges with basic residues in HEAT 19 of Impβ (Fig. 3B). This interaction arrests the helix within the AL of Imp7 near its HEAT 7, that is located on the opposite side of the central cavity, and thus stabilizing Imp7 overall conformation. The second region, Region B, centers on D897 of the AL and its connections to three inter-HEAT loops between HEATs 9-12 of Imp7 and the inter-HEAT loop of HEAT 11of Impβ. The AL emerges from between HEAT 19 helices and is anchored at two additional points, arranging it into a circular shape that blocks the central channel of the heterodimer superhelix. This arrangement enables the binding of the globular domain of H1. The third region, Region C, involves the C-terminal end of the elongated B-helix in HEAT 7 of Impβ which interacts with the elongated loop leading to the subsequent helix (8A). This region contacts the module formed by HEATs 18-19 of Imp7 and the NlB region on the outer surface of Impβ.

All three regions are in close proximity to H1. The H1 globular core domain localizes to the RanGTP-binding domain of Imp7, interacting with the B-helices of HEATs 2–8, while the Lys-rich flexible regions of H1 associate with the Imp7 acidic loop and Impβ, consistent with cross-linking as well as the AF3 predictions (Supplementary Fig. S7).

The fourth interaction region, Region D, involves the structure based re-defined NlB region of Imp7 (residues 1015–1038), which binds within the hydrophobic grooves of HEAT repeats 5-8 in Impβ. On Imp7, the NlB region is situated in close proximity to Region C. Its terminal 15-residue segment (1016–1030) (Fig. 3A) was shown to harbor essential residues for the stable interaction between Imp7 and Impβ ^37^. Specifically, the first conserved GGY/FxF motif of NlB interacts with a hydrophobic groove formed between the A-helices of HEATs 7-8. The VP residues within the second VPyyFzFGG/S motif span helix 7A, extending into the groove between HEAT repeats 6-7, which accommodates the second FxFG motif. Additionally, the C-terminal residues of Imp7 make weak contacts with HEAT 7 and the additional helix of Imp7 (Fig 3A), contributing to the overall stabilization of the complex.

### Biochemical characterization of NIB region binding to Impβ

Despite the remarkable accuracy of AF3 predictions, the limited structural details discernible in the published 6.2 Å resolution cryo-EM map (EMD-0366 ^37^) necessitated additional validation of the proposed structural model. To substantiate the predicted interactions, we characterized the binding of the NlB region to Impβ in greater detail using isothermal titration calorimetry (ITC). The fragment used for this analysis spans residues 1002-1038 of Imp7, previously identified as both necessary and sufficient for Impβ binding ^36^ (Supplementary Fig. S2). Furthermore, ITC measurements were conducted on a series of GST-tagged C– and N-terminally truncated variants of this fragment to delineate the minimal binding region (Table 1, Supplementary Fig. S10). The shortest fragment retaining detectable binding, albeit with approximately fourfold reduced affinity, encompassed residues 1016-1030 of the NlB region. This result aligns exceptionally well with the revised Impβ:Imp7:H1.0-hs structural model (Fig. 3A), which reveals that the NlB fragment from Tyr-1017 to Phe-1030 forms core interactions with Impβ. Pull-down experiments employing deletion mutants further supported these findings (Supplementary Fig. S11). Sequence alignment of homologous importins identified a unique arrangement of conserved residues within the NlB region, including a GG linker (Gly-1015 and Gly-1016) connecting it to the last helix of Imp7. Additionally, the presence of three phenylalanines suggests a role in receptor binding, reminiscent of FG repeats motifs previously implicated in nuclear pore complex studies^37^. Conservation analysis additionally revealed an aromatic residue (Y/F) immediately following the GG linker, giving rise to motifs such as GGY/FxF and VPyyFzFGG/S (Supplementary Fig. S2). Here, “x” represents any amino acid, “y” is primarily Ser or Thr, and “z” corresponds to either Asn or Lys. Interestingly, the aromatic residue Y within the GGY/FxF motif has been shown to be critical for establishing the selectivity barrier in peroxisomes ^55^, while similar FxFG and GFxF motifs have been implicated in nuclear transport receptor binding and stabilization of the nuclear pore complex selectivity barrier ^9^.

**Table 1:** The minimal NlB-region requires residues 1016-1030 of Imp7. The average experimental results (N) are presented, with correlated plots available in Supplementary Figure S12. Highlighted in gray are the non-binders, and dark gray very weak binders.

| Deletions |  |  |  |  |  |  |  |  |  |
| --- | --- | --- | --- | --- | --- | --- | --- | --- | --- |
|  | N | KD [μM] | SD (μM) | ΔH | SD (kJ/mol) | TΔS | SD (kJ/mol) | ΔG | SD (kJ/mol) |
| 1002-1038 | 19 | 0,3 | 0,2 | -69,6 | 15,9 | 22,5 | 2,8 | -37 | 1,4 |
| 1002-1026 | 5 | no binding |  |  |  |  |  |  |  |
| 1002-1029 | 1 | no binding |  |  |  |  |  |  |  |
| 1015-1029 | 3 | no binding |  |  |  |  |  |  |  |
| 1015-1030 | 3 | 1,81 | 0,32 | -62,1 | 5,1 | 29,7 | 5,4 | -32,2 | 0,5 |
| 1015-1033 | 11 | 1,1 | 1,2 | -75,6 | 25,7 | 41,2 | 26,2 | -34,4 | 2,3 |
| 1016-1030 | 2 | 1,41 | 0,1 | -66,9 | 5,6 | 34 | 5,4 | -32,9 | 0,2 |
| 1017-1029 | 3 | no binding |  |  |  |  |  |  |  |
| 1017-1030 | 6 | 315 | 474 | -155,13 | 97,16 | 131,3 | 97,3 | -24,02 | 5,5 |
| 1020-1033 | 2 | no binding |  |  |  |  |  |  |  |
| 1020-1038 | 3 | no binding |  |  |  |  |  |  |  |
| 1025-1033 | 4 | no binding |  |  |  |  |  |  |  |
| 1025-1038 | 5 | no binding |  |  |  |  |  |  |  |

**Table 2:** The effect on various point mutations within the conserved residues of the NlB region of Imp7 are shown, emphasizing the importance of the aromatic residues. The averaged experimental results (N) are presented, with correlated plots available in Supplementary Figure S12A–D.

|  | N | KD [μM] | SD (μM) | ΔH | SD (kJ/mol) | TΔS | SD (kJ/mol) | ΔG | SD (kJ/mol) |
| --- | --- | --- | --- | --- | --- | --- | --- | --- | --- |
| 1002-1038 | 19 | 0,3 | 0,2 | -69,6 | 15,9 | 22,5 | 2,8 | -37 | 1,4 |
| K1009E | 3 | 0,6 | 0,16 | -44,7 | 2,1 | 9,6 | 2,6 | -35 | 0,64 |
| K1013E | 3 | 0,61 | 0,09 | -49,2 | 3,2 | 14,3 | 3,7 | -34,9 | 0,4 |
| G1015P | 1 | 0,19 | n.d. | -58,9 | n.d. | 19,2 | n.d. | -39,8 | n.d. |
| G1015S | 3 | 0,09 | 0,04 | -59 | 2,5 | 19,2 | 3,5 | -39,8 | 1,1 |
| GG15/16SS | 2 | 1,16 | 0,32 | -48,8 | 12,2 | 15,4 | 12,9 | -33,4 | 0,7 |
| GG15/16PP | 2 | 0,4 | 0,07 | -49,6 | 0,3 | 13,8 | 0,7 | -35,9 | 0,6 |
| G1016P | 1 | 0,19 | n.d. | -44,6 | n.d. | 6,88 | n.d. | -37,7 | n.d. |
| G1016S | 4 | 1,23 | 0,32 | -61,1 | 7,8 | 27,9 | 8,4 | -33,2 | 0,7 |
| Y1017S | 2 | no binding |  |  |  |  |  |  |  |
| Y1017G | 2 | no binding |  |  |  |  |  |  |  |
| Y1017F | 6 | 0,3 | 0,08 | -69,5 | 10,8 | 32,8 | 11,2 | -36,7 | 0,7 |
| K1018E | 16 | 1,41 | 0,73 | -59,2 | 20,3 | 37,2 | 27 | -33,2 | 1,1 |
| K1018G | 12 | 1,43 | 0,59 | -70,3 | 27,1 | 37,11 | 17 | -33,2 | 1,1 |
| F1019Y | 16 | 1,36 | 0,32 | -64,7 | 21,1 | 31,6 | 21 | -33 | 0,6 |
| F1019G | 3 | no binding |  |  |  |  |  |  |  |
| F1019S | 4 | no binding |  |  |  |  |  |  |  |
| YF17/19FY | 4 | 3,8 | 1,1 | -68,2 | 16,7 | 37,7 | 16,7 | -30,6 | 0,8 |
| N1020S | 4 | 0,16 | 0,05 | -70,4 | 5,8 | 32,1 | 5,2 | -38,3 | 0,7 |
| V1024E | 3 | 4,66 | 2,18 | -48,8 | 18,8 | 18,6 | 19 | -30,9 | 1,2 |
| V1024T | 10 | 1,11 | 0,6 | -63,9 | 16,4 | 30,14 | 15,5 | -33,7 | 1,3 |
| P1025G | 6 | 2,02 | 1,5 | -63,1 | 14,3 | 30,7 | 4,6 | -32,4 | 1,4 |
| F1028G | 2 | no binding |  |  |  |  |  |  |  |
| F1028S | 11 | no binding |  |  |  |  |  |  |  |
| F1028Y | 4 | 1,72 | 0,88 | -49 | 12,9 | 16,4 | 14,1 | -32,6 | 1,2 |
| N1029S | 5 | 0,17 | 0,03 | -71,2 | 8,9 | 33,2 | 9 | -38 | 0,4 |
| F1030G | 3 | no binding |  |  |  |  |  |  |  |
| F1030S | 8 | no binding |  |  |  |  |  |  |  |
| F1030Y | 8 | 20,9 | 27 | -41 | 16,9 | 14,9 | 16,8 | -28,7 | 4 |
| FF28/30SS | 5 | no binding |  |  |  |  |  |  |  |
| FF28/30YY | 3 | no binding |  |  |  |  |  |  |  |
| G1031P | 3 | 0,31 | 0,14 | -80,4 | 27 | 43,5 | 27,8 | -36,8 | 1,22 |
| G1031I | 5 | 0,44 | 0,07 | -54,5 | 8,4 | 18,7 | 8,7 | -35,7 | 0,4 |
| N1032G | 3 | 0,16 | 0,02 | 82,1 | 9,4 | 44 | 9,2 | -38,1 | 0,3 |

To further validate the arrangement of the NlB region on Impβ, we introduced mutations in the conserved residues of the GST-tagged Imp7 (1002-1038) fragment to assess their impact on the binding affinity and interaction properties via ITC. Single and some double-point mutations were designed to test the structural model and the functional significance of key interaction sites (Supplementary Fig. S12). As anticipated, substitutions of the four conserved key aromatic residues – Y1017, F1019, F1028, and F1030 – with glycine (G) or serine (S) abolished binding, consistent with earlier pull-down experiments^37^. Double mutations (Y1017/F1019 or F1028/F1030 to G or S) had similarly disruptive effects. In contrast, aromatic-to-aromatic substitutions (e.g.: Y-to-F or F-to-Y) revealed position-specific effects: the Y1017F mutation did not affect binding affinity, indicating that the hydroxyl group at this position is dispensable. However, F-to-Y mutations at F1019, F1028, or F1030 caused at least a five-fold reduction in affinity, suggesting that the additional hydroxyl group destabilizes interactions by interfering with the aromatic packing pattern (Fig. 3A). Mutations in the conserved GG linker (G1015– G1016) further highlighted its role in stabilizing the NlB region. Proline substitutions (G1015P, G1016P) enhanced binding affinity by two-to three-fold, whereas the double mutation of both glycines to proline showed no significant change in binding (similar to the wild type protein), suggesting redundancy in their stabilizing role. The G1016S mutation or the double G1015S/G1016S substitution significantly impaired binding, pointing to a critical role of G1016 in enabling the optimal orientation of the NlB region.

Unexpectedly, the conserved asparagine residue N1020 exhibited increased binding affinity upon mutation to serine (N1020S), indicating a degree of structural plasticity at this position. In contrast, the VP motif located between the two FxFG motifs emerged as a critical hydrophobic anchor. Mutation of Val1023 to glutamate (V1023E) or threonine (V1023T) drastically reduced binding affinity (15– and 4-fold, respectively), while Pro1024 to glycine (P1024G) substitution caused a seven-fold reduction in binding, underscoring its role in orienting the NlB region for optimal Impβ interaction (Fig. 3A). Mutations in the downstream residues N1029, G1031, and N1032 further highlighted their functional contributions. While N1029S and N1032G substitutions modestly enhanced binding affinity (∼2-fold), proline or isoleucine substitutions at G1031 had no significant effect, suggesting that flexibility at this position is less critical for interaction.

Overall, ITC measurements strongly support the proposed interaction mode between the NlB region of Imp7 and Impβ, reinforcing the functional relevance of these key residues in mediating stable interactions within the ternary import complex.

## DISCUSSION

The binding interaction between Imp7 and Impβ has been extensively characterized biochemically, demonstrating Imp7’s dual role as both a chaperone and transport receptor within the Impβ:Imp7 heterodimer ^33^. The interaction with H1 is indirectly dependent on the NlB region of Imp7, which facilitates cooperative binding between Imp7 and Impβ, acting as a prerequisite for H1 association and transport^34,36^. To further investigate the allosteric regulation of the Imp7-Impβ interaction, we utilized AF3 to generate models of the heterotrimeric complex comprising Impβ, Imp7 and either of two different histone H1 molecules: H1.11L-chick ^36^ and H1.0-hs^37^. While both histones share the structurally similar core globular domain (H1.11L-chick: region 41–114; H1.0-hs: region 20–97), they differ in the length of their N-terminal fragment. Across multiple independent simulations, a consistent model emerged in which Impβ and Imp7 formed an extended superhelical arrangement, with the core domain of both histones positioned almost identically within Imp7. This arrangement contradicts the 6.2 Å cryo-EM structure of the Imp7:Impβ:H1.0-hs complex^37^ (PDB id: 6N88), which suggested an inverted arrangement, where the histone core domain is bound within Impβ. Remarkably, the cross-linking data (BS3, MTMM)^37^ overlooked during structure determination, align with the predicted Impβ:Imp7:H1.0-hs^AF3^ heterotrimeric complex (Fig. 1, Supplementary Table S4). Similarly, our own BS3 cross-link results support the predicted Impβ:Imp7:H1.11L-chick^AF3^ complex (Supplementary Tables S3, S5, Fig. 1). Refinement of the Impβ:Imp7:H1.0-hs^AF3^ model against the available cryoEM map (EMD-0366) yielded a revised structure of the Impβ:Imp7:H1.0-hs complex. Validation statistics (Fig. 2, Supplementary Table S6) strongly support this refined model. Nevertheless, due to the limited resolution (6.2 Å) of the cryo-EM map additional validation was performed with various mutants using ITC measurements and pull-down experiments.

In the Imp7:Impβ:H1.0-hs complex (PDB ID: 6N88), Impβ adopts an overall extended conformation with highly rotated terminal HEAT repeats, likely forced to fit the cryo-EM map corresponding to significantly larger Imp7, resulting in a register shift. However, no such distortions are observed when Imp7 occupies this position (Supplementary Fig. S6, S9). The C-terminal domain of Imp7, comprising the last five helices, forms a stable structural entity that fits well into the map, bridging the two molecules while maintaining its interaction with Imp7’s HEAT 7B helix. In contrast, the deposited 6N88 structure requires significant rearrangement of the C-terminal HEAT repeats of Impβ to accommodate the map – an unprecedented conformation in any previously determined Impβ structures (Supplementary Fig. S6, S9). This discrepancy strongly suggests the spatial arrangement of both importins within the complex was inverted, thereby hindering a precise molecular understanding of the ternary complex. The revised Impβ:Imp7:H1.0-hs structural model provides novel insights into the functional roles of the acidic loop (AL) and the NlB region of Imp7 in the ternary complex formation. Independent validation by ITC and pull-down experiments confirmed that the key residues required for stable interaction with Impβ are located within the 1016-1030 region of Imp7’s NlB. This finding aligns with the structural data, as the preceding residues form part of the elongated C-terminal α-helix of Imp7 (Fig. 3A), which, together with the terminal two HEAT repeats, constitutes a stable interaction module. This module is crucial for binding the unusually extended B-helix of HEAT 7 and its adjacent loop residues (Fig. 3C). Within the 1016-1030 region, the phenylalanine residues embedded in two conserved motifs of the NlB, GYxF and FxFG, bind to the convex outer surface of Impβ, forming hydrophobic interactions reminiscent of those observed between karyopherins and FG-repeat nucleoporins ^37^. FG motifs are critical elements of the NPCs selectivity barrier, serving as transient interaction sites with karyopherins during facilitated translocation^56–63^. Structural studies have consistently mapped FG-binding sites to the convex outer surface of nuclear transport receptors (NTRs), specifically within the grooves formed by A-helices of adjacent HEAT repeats ^60,64–67^; see also Supplementary Fig. S13). This outer-surface localization ensures that FG-binding does not interfere with cargo or RanGTP interactions, which predominantly occur along the concave inner surface formed by the B-helices. In Impβ, multiple hydrophobic FG-binding sites have been experimentally identified, with additional binding sites suggested by molecular dynamics (MD) simulations (Supplementary Fig. S13). These findings reinforce the hypothesis that the NlB region binds Impβ in a manner analogous to FG-repeat nucleoporins, highlighting the evolutionary conservation of FG-mediated interactions in nucleocytoplasmic transport.

Two major binding modes for FG-Nups have been structurally characterized in members of the Impβ superfamily. In the first, more complex binding mode, a phenylalanine (F) residue from the FXF motif inserts into an aromatic cavity, where π-π stacking interactions with an adjacent aromatic residue stabilize the interaction. The second phenylalanine of the motif further shields this binding site from solvent exposure through additional π-stacking interactions. The second, simpler mode involves hydrophobic or aromatic interactions between a single phenylalanine residue and a surrounding cleft. Structural analyses reveal that the NlB region engages Impβ predominantly through the first binding mode for both conserved sequence motifs: GYxF and FxFG (Fig. 3), highlighting the mechanistic parallels between the NlB region and FG-Nups. This mode of interaction exemplifies the adaptive plasticity of importins in recognizing FG motifs across diverse binding partners. ITC experiments using point mutations in aromatic residues further corroborate this interaction model, placing the NlB region within the hydrophobic clefts formed by the A-helices of Impβ HEAT repeats 6–8.

These results align with previous deletion mutant studies ^34,36^. Within the first conserved motif (GGYxF), F1019 is deeply embedded in the cleft formed by HEATs 7 and 8, forming key hydrophobic interactions, while the upstream aromatic residue (Y) acts as a shielding cap, a feature, reflected in sequence alignments across species (alignment, Supplementary Fig. S2). All other mutations tested at this site abolished binding, highlighting the specificity of this interaction. In the second FxFG motif, both phenylalanines are inserted into the cleft between HEAT repeats 6 and 7 of Impβ, consistent with a previously characterized FG-Nup binding mechanisms ^66^. The five C-terminal residues of the NlB region exhibit considerable flexibility and are dispensable for binding, as their deletion has little impact on affinity. Importantly, mutating six residues within the HEAT 5-6 and HEAT 6-7 clefts (L174S, T175A, I178D, E214A, F217A, and I218D) significantly reduces or abolishes Imp7 binding ^37^. According to our model, the mutations in the HEAT 6-7 cleft (E214A, F217A, and I218D) exert the strongest impact. These findings emphasize the dominant role of the HEAT 6-7 binding groove in stabilizing the Imp7-Impβ interaction ^37^.

The revised Impβ-Imp7-H1 structural model aligns closely with available biochemical data, offering mechanistic insights into the distinct binding affinities of individual transport receptors for H1. While full-length H1 exhibits weak binding affinity for Impβ, it binds moderately to Imp7^34^ and displays stronger preference for Imp7 over Impβ in ITC measurements^36^. Consistent with these observations, the heterodimeric Impβ:Imp7 complex has been demonstrated to simultaneously bind both H1 and the IBB-domain of Impα^34^. The revised structural model supports this observation, as the IBBα domain can be positioned within the complex without interfering with H1 binding to Imp7 (Supplementary Fig. S14).

Previous findings supporting the placement of H1 globular core within the concave surface of Impβ include stronger H1-core binding to Impβ in pull-down assays and its nuclear import via Impβ in an *in vitro* assay^35^. Structurally, the globular domain of H1 contains multiple surface-exposed basic residues, which could facilitate electrostatic interactions with the acidic concave interior of Impβ. However, comparison with the crystal structure of the hsSREBP bound to mouse Impβ (PDB ID: 1UKL), where a dimeric globular domain of SREBP occupies the position originally assigned to the H1 core domain in the Imp7:Impβ:H1hs complex, suggests that such binding might be an artifact. In the context of full-length H1, the remaining unstructured regions are likely to sterically hinder these interactions, further supporting the revised model in which H1 core domain binds exclusively to Imp7. The burial or occlusion of critical residues in the intact H1 molecule may further impede direct interaction with the concave surface of Impβ (Supplementary Fig. S15).

### Proposed model for assembly and disassembly of the transport complex

Biochemical analyses provide insights into the assembly mechanism of the Imp7:Impβ:H1 complex. These experiments demonstrate that preassembled Impβ-H1 complexes undergo a stepwise transition into an import-competent state upon Imp7 addition. In contrast, no such transition occurs when Imp7 first associates with H1, suggesting that the NIB region cannot effectively engage Impβ once H1 is bound to Imp7 ^35,36^. This observation suggests that Imp7:Impβ complex formation proceeds through a multistep process. Initially, the NlB region of Imp7 binds to Impβ, stabilizing the heterodimeric complex – consistent with the lack of detectable interaction between Imp7 and Impβ when the NlB region is deleted ^35^.

Subsequent steps in the assembly of the Imp7:Impβ:H1 complex are likely driven by the interaction of Impβ with the C-terminal module of Imp7 (region C). This interaction is preceded by histone H1 integration, which induces an increase in the inner diameter of Imp7 and a pronounced change in curvature, enabling the formation of region C. Binding of H1 to Imp7 also triggers conformational changes in the extended 74-residue acidic loop (AL), positioning the C-terminal α-helix of H1 between Imp7 and Impβ. The AL undergoes structural rearrangement, with a helical fragment bridging Imp7 and Impβ, thereby facilitating the formation of interaction region B and subsequently A. Surprisingly, deletion of the last 258 residues of Impβ, which disrupts region A formation, does not impair nuclear import of H1, indicating that this region plays an auxiliary rather than essential role in the nuclear translocation process ^34^.

The transport-competent import complex passively traverses the permeability barrier of the nuclear pore complex (NPC). The occupation of two prominent FxFG binding sites on Impβ by the NlB region of Imp7 may modulate Impβ’s binding properties and specificity. This could impact the transport pathway by excluding certain nucleoporins from binding, accelerating the passage through the NPC, or altering the site of complex disassembly. Upon reaching the nuclear basket, where Nup153 is located, the complex is retained, as Nup153 has been shown to bind strongly to Impβ ^60,61^. The binding of RanGTP subsequently terminates the import process by triggering structural rearrangements in Impβ. These changes reduce Impβ’s affinity for Imp7, displacing the NlB region from its binding site and releasing the Imp7:H1 complex into the nucleoplasm. Once released, the Imp7:H1 complex may interact with chaperones such as Nuclear Autoantigenic Sperm Protein (NASP) ^68^. NASP, which forms a dimer and contains a high proportion of acidic residues (25%; UniProt ID: P49321), could neutralize the highly basic nature of H1 ^68^. Binding of the basic C-terminal tail of H1 to NASP may facilitate H1 release from Imp7, while promoting RanGTP binding to Imp7. This step dissociates the import complex, allowing Imp7 to be recycled back to the cytoplasm. This putative mechanism resembles the transport pathway of monomeric histone H3 by Imp5, in which NASP acts as a nuclear chaperone upon histone release^46^. Imp7:H1-bound NASP, possibly in coordination with additional histone chaperones, may guide H1 to assembling nucleosomes, ensuring its proper deposition onto DNA and histone octamers.

## OUTLOOK

The use of the AI prediction tool AlphaFold3, provided an unprecedentedly reliable model of the ternary Impβ:Imp7:H1 complex. This model is strongly supported by cross-linking data and ITC measurements. Moreover, the predicted structure demonstrated excellent agreement with the cryoEM map, significantly revising its initial structural interpretation.

Thus, while AlphaFold3 generates highly accurate initial models, experimental validation remains essential to refine and confirm structural details. Integrating predictive modeling with experimental approaches, such as cryo-EM, cross-linking mass spectrometry, and thermodynamic binding assays, ensures a more precise understanding of macromolecular complexes. Future studies combining these techniques will further enhance our knowledge of nuclear transport mechanisms and provide deeper insights into the conformational dynamics of transport receptors and their cargos.

## RESOURCE AVAILABILITY

### Data and code availability

– All data reported in this paper have been deposited to the PDB (9QEJ and 9QF0).
– The results of the crosslinking experiments can be found in the Supplementary data section.
– This work did not generate any new code.

## ACKNOWLEDGEMENTS

We thank A. Berndt and J. Heine for technical assistance and many generations of Bachelor students for performing ITC experiments and R. Kehlenbach for carefully and critically reading the manuscript. RF and HU were supported by the Deutsche Forschungsgemeinschaft (SFB860 and SFB 1565 (project number 469281184).

## AUTHOR CONTRIBUTIONS

O.D. performed, O.D. and H.U. analyzed the crosslinking data. P.N. performed modelling, structure determination and structure – crosslinking interpretation. A.D. performed protein expression and purification, ITC measurements, and pulldown experiments. P.N, R.F. and A.D. conceptualized and supervised the study. All authors reviewed and edited the manuscript. R.F. managed project administration and funding acquisition.

## STAR*METHODS

Detailed methods are provided in the online version of this paper and include the following:

〇 KEY RESOURCES TABLE
〇 METHOD DETAILS
  〇 **Cloning of NlB-Fragments**
  〇 **Recombinant protein expression and protein purification**
    ■ **Expression**
    ■ **Purification**
  〇 **Pull down experiments**
  〇 **ITC: NlB – Importin β Interaction by Isothermal Calorimetry**
  〇 **Identification of protein crosslinks by Mass Spectrometry**
  〇 **Prediction of complexes and their validation using cross-link data**
  〇 **Re-evaluation of Cryo-EM structures: Imp7:Impβ:Histone-H1.0 and Impβ:Histone-H1.0**

## SUPPLEMENTAL INFORMATION

Supplemental information can be found online at the end of this manuscript.

## Supplemental Information

## STAR*METHODS

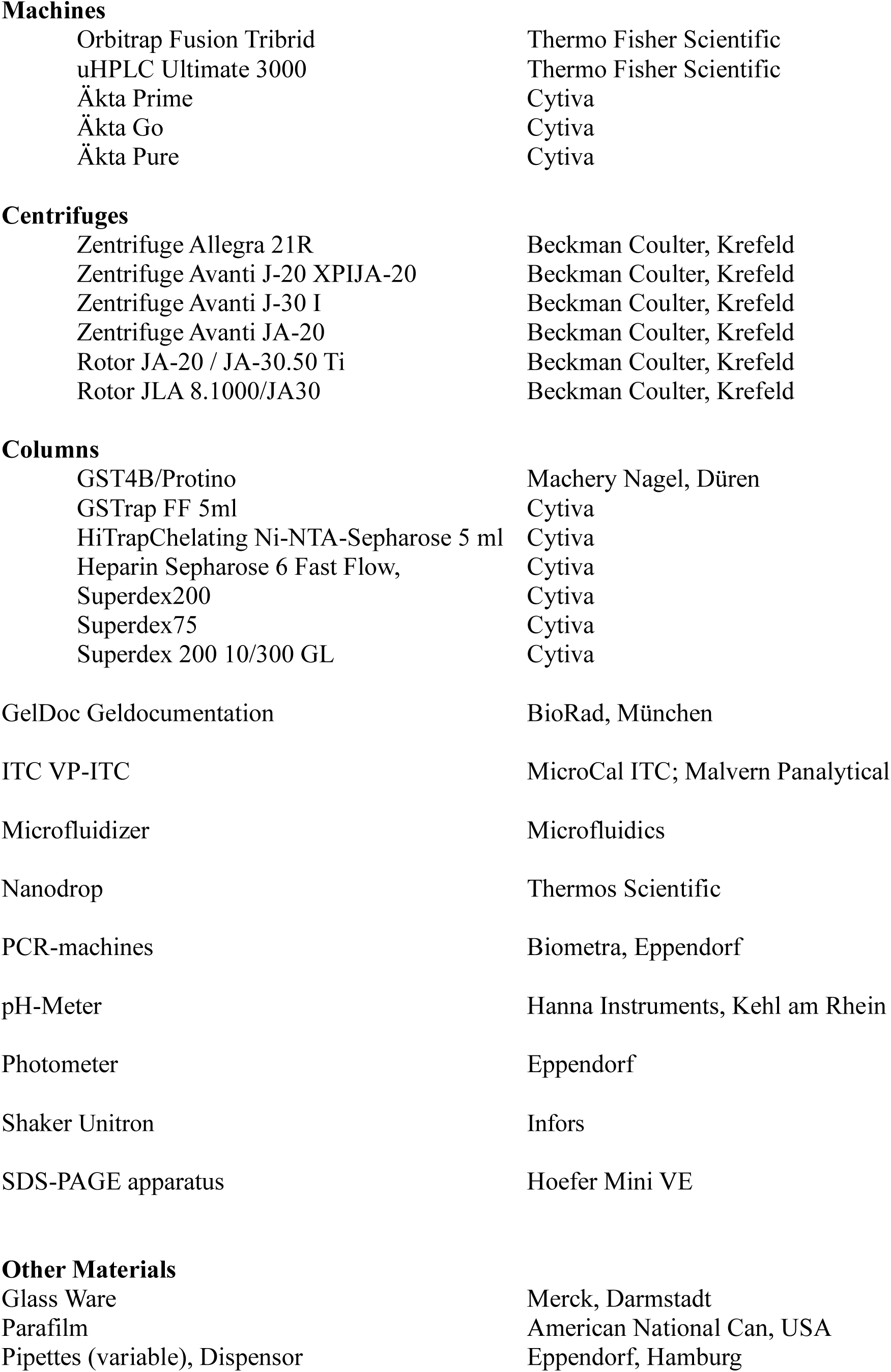
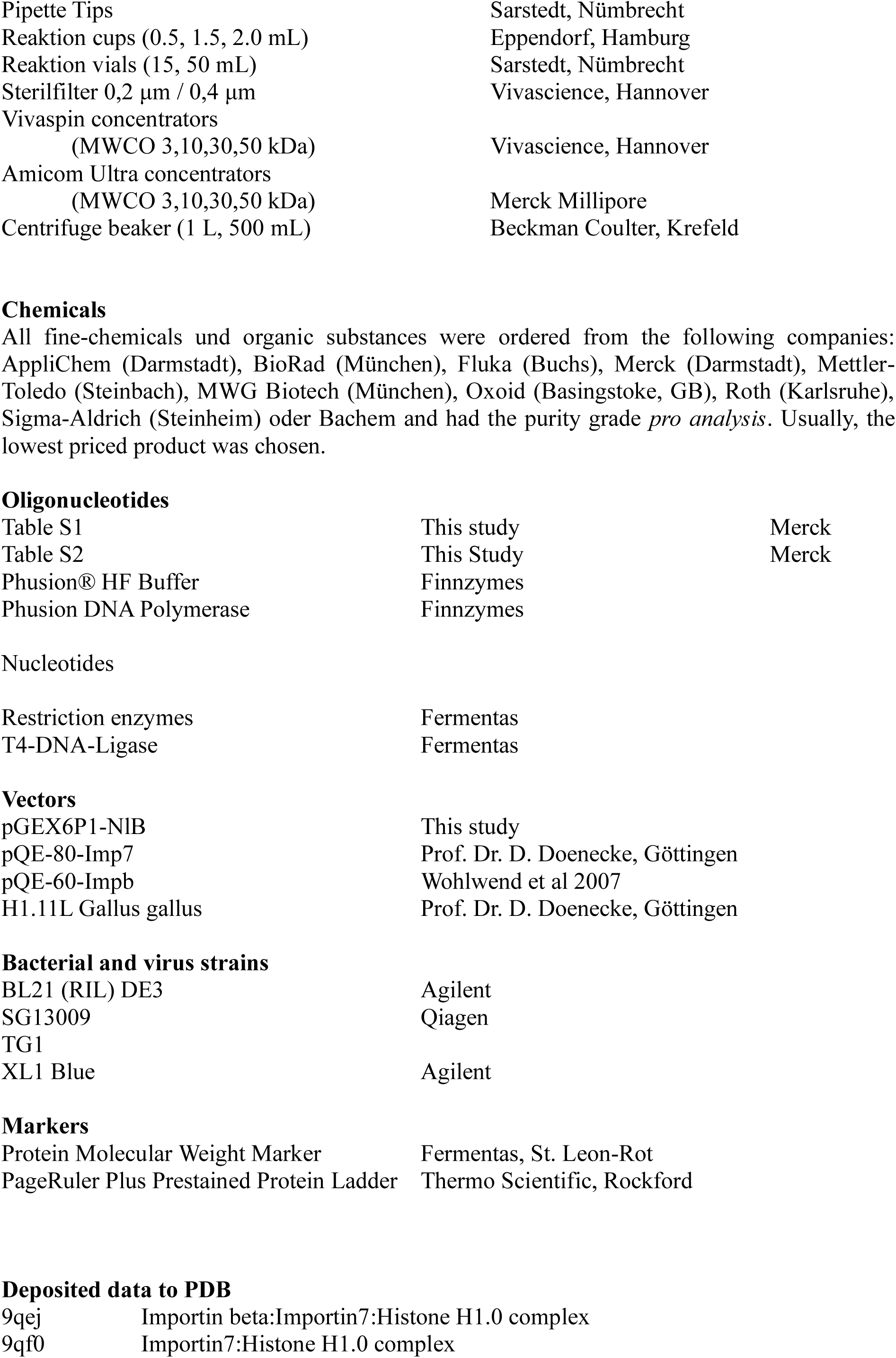
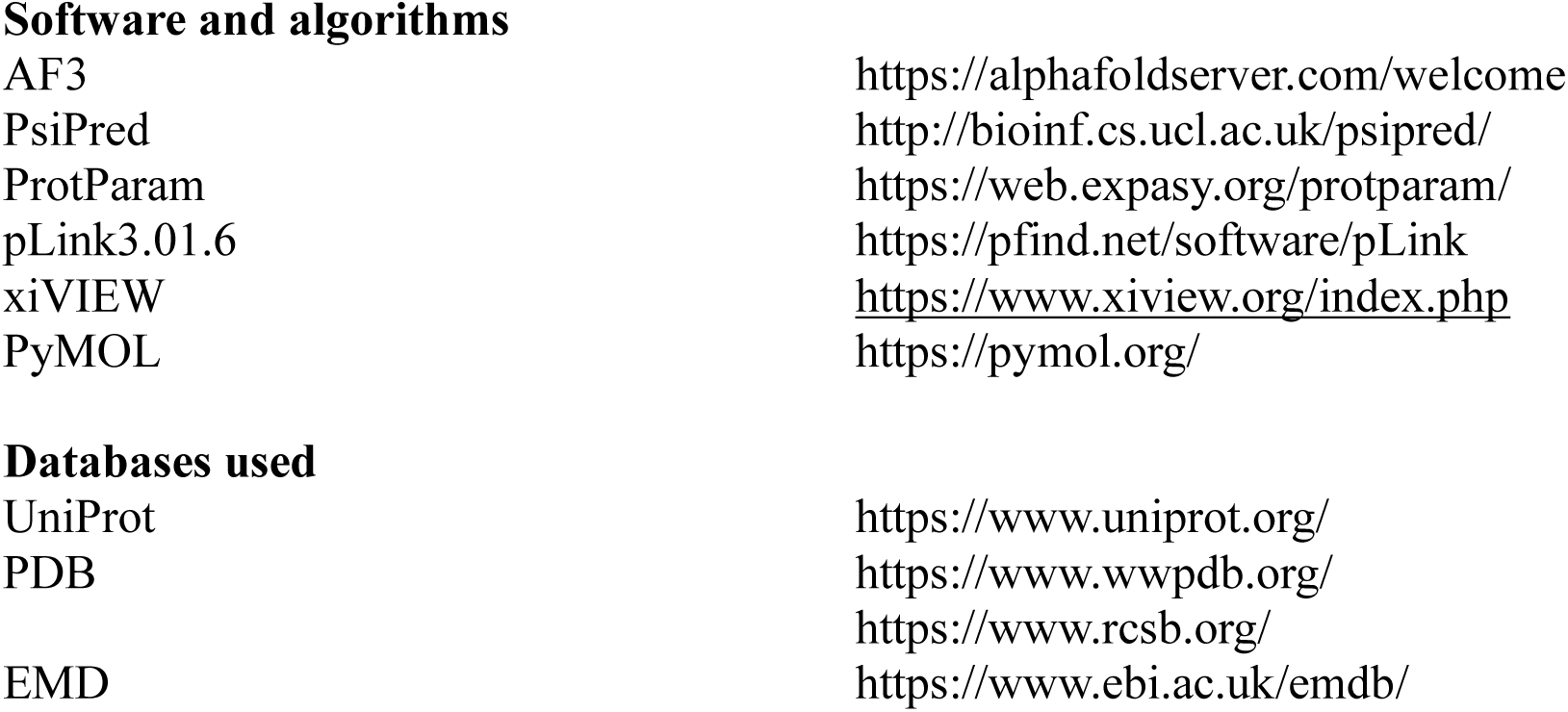
KEY RESOURCES TABLE.

Machines
|  |  |
| --- | --- |
| Orbitrap Fusion Tribrid | Thermo Fisher Scientific |
| uHPLC Ultimate 3000 | Thermo Fisher Scientific |
| Äkta Prime | Cytiva |
| Äkta Go | Cytiva |
| Äkta Pure | Cytiva |
| Centrifuges |  |
| Zentrifuge Allegra 21R | Beckman Coulter, Krefeld |
| Zentrifuge Avanti J-20 XPIJA-20 | Beckman Coulter, Krefeld |
| Zentrifuge Avanti J-30 I | Beckman Coulter, Krefeld |
| Zentrifuge Avanti JA-20 | Beckman Coulter, Krefeld |
| Rotor JA-20 / JA-30.50 Ti | Beckman Coulter, Krefeld |
| Rotor JLA 8.1000/JA30 | Beckman Coulter, Krefeld |
| Columns |  |
| GST4B/Protino | Machery Nagel, Düren |
| GSTrap FF 5ml | Cytiva |
| HiTrapChelating Ni-NTA-Sepharose 5 ml | Cytiva |
| Heparin Sepharose 6 Fast Flow, | Cytiva |
| Superdex200 | Cytiva |
| Superdex75 | Cytiva |
| Superdex 200 10/300 GL | Cytiva |
| Other Materials |  |
| Glass Ware | Merck, Darmstadt |
| Parafilm | American National Can, USA |
| Pipettes (variable), Dispensor | Eppendorf, Hamburg |
| Pipette Tips | Sarstedt, Nümbrecht |
| Reaktion cups (0.5, 1.5, 2.0 mL) | Eppendorf, Hamburg |
| Reaktion vials (15, 50 mL) | Sarstedt, Nümbrecht |
| Sterilfilter 0,2 µm / 0,4 µm | Vivascience, Hannover |
| Vivaspin concentrators<br>(MWCO 3,10,30,50 kDa) | Vivascience, Hannover |
| Amicom Ultra concentrators<br>(MWCO 3,10,30,50 kDa) | Merck Millipore |
| Centrifuge beaker (1 L, 500 mL) | Beckman Coulter, Krefeld |

Oligonucleotides
|  |  |  |
| --- | --- | --- |
| Table S1 | This study | Merck |
| Table S2 | This Study | Merck |
| Phusion® HF Buffer | Finnzymes |  |
| Phusion DNA Polymerase | Finnzymes |  |
| Nucleotides |  |
| Restriction enzymes | Fermentas |
| T4-DNA-Ligase | Fermentas |
| Vectors |  |
| pGEX6P1-NIB | This study |
| pQE-80-Imp7 | Prof. Dr. D. Doenecke, Göttingen |
| pQE-60-Impb | Wohlwend et al 2007 |
| H1.11L Gallus gallus | Prof. Dr. D. Doenecke, Göttingen |
| Bacterial and virus strains |  |
| BL21 (RIL) DE3 | Agilent |
| SG13009 | Qiagen |
| TG1 |  |
| XL1 Blue | Agilent |
| Markers |  |
| Protein Molecular Weight Marker | Fermentas, St. Leon-Rot |
| PageRuler Plus Prestained Protein Ladder | Thermo Scientific, Rockford |
| Deposited data to PDB |  |
| 9qej | Importin beta:Importin7:Histone H1.0 complex |
| 9qf0 | Importin7:Histone H1.0 complex |
| Software and algorithms |  |
| AF3 | https://alphafoldserver.com/welcome |
| PsiPred | http://bioinf.cs.ucl.ac.uk/psipred/ |
| ProtParam | https://web.expasy.org/protparam/ |
| pLink3.01.6 | https://pfind.net/software/pLink |
| xiVIEW | https://www.xiview.org/index.php |
| PyMOL | https://pymol.org/ |
| Databases used |  |
| UniProt | https://www.uniprot.org/ |
| PDB | https://www.wwpdb.org/ |
|  | https://www.rcsb.org/ |
| EMD | https://www.ebi.ac.uk/emdb/ |

## METHOD DETAILS

### Cloning of NlB-Fragments

Deletion constructs of the *X. laevis* NlB-region (Supplementary Table S1) were cloned into the pGEX-6P1 vector using BamHI and XhoI restriction sites. The fragments were amplified via standard PCR from a full-length NlB region clone (Imp7 residues 1002-1038), digested with the selected restriction enzymes, gel-purified, and ligated into the double-digested, dephosphorylated vector following standard molecular cloning methods.

Short DNA fragments corresponding to the NlB region were introduced by primer annealing and subsequent ligation into BamHI– and XhoI-cleaved pGEX-6P1. Point mutations were designed based on sequence alignments of homologous proteins from various organisms (Supplementary Fig. S2). These mutations were generated using QuickChange PCR, followed by DpnI digestion to remove the template DNA. Primers used for PCR amplifications are listed in Supplementary Table S2.

A linker insertion, introducing the additional amino acid sequence GSGGGSGGGTGGGSG before the NlB sequence, was performed by primer annealing and ligation into the BamHI site of the respective clones. All cloned variations of the NlB region were confirmed by colony picking, plasmid DNA amplification, isolation, and sequencing.

### Recombinant protein expression and protein purification

#### Expression

Human Importin β and *X. laevis* Imp7 NlB region, along with its variants, were expressed and purified using the following protocol. The *H. sapiens* pQE60-Impβ construct was transformed into *E. coli* SG13009 cells. An overnight pre-culture was used to inoculate 500 mL of 2YT medium at a 1:50 ratio. The cells were grown at 37 °C until reaching an OD_600_ of 1.2, then diluted 1:2 with pre-cooled (4 °C) medium. Expression was induced by adding IPTG to a final concentration of 300 µM, followed by the addition of K₂HPO₄ to a final concentration of 30 mM. The culture was incubated overnight at 16 °C. The cells were harvested by centrifugation at 4800 × g, washed once with PBS, frozen in liquid nitrogen, and stored at −20 °C until further use. The Imp7 NlB region and its variants were cloned into pGEX-6P1 and expressed as GST-fusion proteins. After transformation into *E. coli* BL21(DE3) cells, the expression protocol was identical to that described for Importin β, with the exception that IPTG was added to a final concentration of 250 µM, and K₂HPO₄ was omitted.

#### Purification

The cell pellets were resuspended in the respective lysis buffers (His-tagged Impβ: 50 mM Tris/HCl pH 7,5, 200 mM NaCl, 10 mM Imidazole; GST-fusions: 50 mM Tris/HCl pH 7,5, 100 mM NaCl, 2 mM DTT) and the cells disrupted using a pneumatic disintegrator (Microfluidizer 110S, Microfluidics). The lysate was cleared by centrifugation at 20,000 × g for 30 min. Both proteins were purified in a two-step process. His-tagged Impβ was purified using a His60 Ni Superflow^TM^ (Clontech) or HisTrap^TM^ FF columns (Cytiva). After loading of the sample, the column was washed and Impβ eluted using a step gradient up to 500 mM Imidazole in lysis buffer. GST-NlB versions were purified by applying the supernatant on a GSH-Sepharose column and after a washing step eluted with a buffer composed of 50 mM Tris Base, 100 mM NaCl, 30 mM reduced Glutathione.

The second purification step for all proteins was a size exclusion chromatography using either a Superdex 75 16/60 or a Superdex 200 16/60 (Cytiva) equilibrated in 20 mM Tris/HCl pH 7.5, 100 mM NaCl at a flow rate of 1 mL/min. Histone H1.11L from chicken was prepared as described previously ^1^. See supplemental Figure S3 for comparison to H1 from human.

#### Pull down experiments

Approximately 10–20 nmol of GST-NlB fragments, deletions or point mutations were coupled to GSH-Sepharose 4B beads (Cytiva) per experiment, equilibrated in 20 mM Tris/HCl pH 7.5, 100 mM NaCl. After incubation for 30 min at 20°C with shaking at 600 rpm in a thermostat, the beads were pelleted by centrifugation at 500 × g for 5 min.

The beads were washed three times with 1 mL of buffer. For each washing step, buffer was added, the beads were resuspended, centrifuged at 500 × g for 5 min at 20°C, and the supernatant was removed. After the final wash, any remaining liquid was carefully removed from the beads, and 80 µL of water was added. The sample was vigorously vortexed before adding 80 µL of 2× Laemmli buffer.

The samples were boiled at 95°C for 10 min, centrifuged at 500 × g for 5 min at 20°C, and then applied to a 12.5% SDS-PAGE gel. Gels were subsequently stained using Coomassie solution.

#### ITC: NlB – Importin β Interaction by Isothermal Calorimetry

All ITC experiments were performed using a VP-ITC microcalorimeter (Malvern, originally MicroCal Inc.; reaction cell volume: 1462 µL). His-tagged Impβ and GST-tagged NlB region variants were purified as described above, ensuring the use of the same buffer (20 mM Tris pH 7.5, 100 mM NaCl, 2mM MgCl_2_) during the final purification step via size exclusion chromatography before ITC measurements. In a typical setup, Impβ (5–12 µM) was placed in the sample cell, while the ligand (GST-NlB or its fragments, 70–200 µM) was loaded into the titration syringe. Experiments were conducted at 20°C with a reference power of 10–12 and a stirring speed of 507 rpm. The titration protocol consisted of one initial injection of 4 µL (over 8 seconds), followed by 18 injections of 15 µL (over 30 seconds), with an interval time of 300 – 360 s between injections. Data analysis was performed using MicroCal PEAK-ITC analysis software, with the complex stoichiometry preset to N = 1. Protein concentrations were determined using either the standard Bradford reagent (Rothi-Quant, Carl Roth), the calculated extinction coefficient ε_280_ (Expasy: ProtParam, http://web.expasy.org/protparam/) using a NanoDrop, or the method described by Ehresmann and Weil ^2^.

#### Identification of protein crosslinks by Mass Spectrometry

One hundred fifty pmol of the purified trimeric complex Impβ:Imp7:H1.11L-chick was crosslinked with 150 µM BS3 in a volume of 50 µl for 30 min at 24 °C and subsequently quenched with 20 mM Tris-HCl, pH 8. Fifty pmol of the crosslinked sample were resolved on a 4-12% Bis-Tris NuPAGE gel (Thermo Fisher Scientific), protein bands were visualized by Coomassie staining, the band of the crosslinked trimeric complex was excised, proteins were reduced with DTT, alkylated with iodoacetamide and in-gel digested with trypsin (Promega). Extracted peptides were dried in a SpeedVac vacuum concentrator (Thermo Fisher Scientific), dissolved in 25 µl 5% acetonitrile / 0.1% trifluoroacetic acid and analyzed by uHPLC-MS. For this, peptides were on-line separated on a custom C18 reverse-phase self-packed column (ReproSil-Pur 120 C18-AQ, 1.9 µm pore size, 75 µm inner diameter, 30 cm length; Dr. Maisch GmbH) using an Ultimate 3000 uHPLC coupled to Orbitrap Fusion Tribrid Mass Spectrometer (both Thermo Fisher Scientific) with a Nanospray Flex Ion source. Data were acquired using a 60 or 90 min method employing either a TopN approach (selecting the 20 most abundant precursors) or Top Speed strategy with a 3 s data-dependent acquisition cycle. One full MS scan across the 350–1600 m/z range was acquired at a resolution of 120000, with an AGC target of 5E+5 and a maximum fill time of 100 ms. Precursors with charge states 3–8 above a 5e4 intensity threshold were then sequentially selected using isolation window of 1.6 m/z, subjected to higher-energy collisional dissociation at a normalized collision energy setting of 30% (HCD), and the resulting MS2 spectra were recorded at a resolution of 30000, AGC targets of 5E+4 and a maximum fill time of 128 ms. Dynamic exclusion of precursors was set to 10 or 20s. BS3-mediated protein-protein crosslinks were identified by pLink3.0.16 ^3^ by searching Thermo raw files against a database containing 3 target proteins and trypsin sequences. Carbamidomethylation of cysteine was set as a fixed modification, acetylation of protein N-terminus and oxidation of methionine as variable. The results were filtered to a false discovery rate of 5% at peptide level.

#### Prediction of complexes and their validation using cross-link data

Atomic models of Impβ:Imp7:H1.11L-chick and Impβ:Imp7:H1.0-hs complexes were generated using the AlphaFold3 (AF3) server (https://alphafoldserver.com/)^4,5^ for the full-length sequences of *Xenopus laevis* Importin7 (Imp7), *Homo sapiens* Importinβ (Impβ), and histones: *Gallus gallus* H1.11L (chicken) and *Homo sapiens* H1.0. To enhance structural diversity in predictions, 20 individual runs with differing random seeds were submitted for the Impβ:Imp7:H1.11L-chick complex, resulting in 100 predictions (Supplementary Fig. S7). In contrast, only 5 predictions were obtained for the Impβ:Imp7:H1.0-hs complex. All atomic models were superposed onto a randomly selected reference Impβ:Imp7:H1.11L-chick complex, aligning the Impβ and Imp7 molecules, which are identical in both complexes, using PyMol. RMSDs were calculated for all atoms. To assess differences in the positioning of histone molecules within the superposed models, RMSD calculations were performed relative to the histone molecule in the reference one, focusing specifically on the core globular domain (H1.11L-chick: region 41–114; H1.0-hs: region 20–97, Fig. 1). For these RMSD calculations, no additional structural alignment was applied, employing the PyMOL “align” command with the options: “cycles=0, transform=0, object=aln”. All predicted models, 100 Impβ:Imp7:H1.11L-chick and 5 Impβ:Imp7:H1.0-hs complexes, were checked for their compatibility with available cross-link data using a custom bash/awk script. This script measured distances between Cα atoms of cross-linked amino acid residues and categorized them based on defined thresholds for BS3 cross-linker: distances < 24 Å were classified as optimally fulfilled, < 29 Å as generously fulfilled, and > 29 Å as not fulfilled (Fig. 1, Supplementary Tables S4, S5).

#### Re-evaluation of Cryo-EM structures: Imp7:Impβ:Histone-H1.0 and Impβ:Histone-H1.0

The atomic models and corresponding cryo-EM maps of the Imp7:Impβ:Histone-H1.0 and Impβ:Histone-H1.0 complexes were obtained from the Protein Data Bank (PDB). The Imp7:Impβ:Histone-H1.0 complex was deposited under the accession code 6N88, with the associated cryo-EM map at a reported resolution of 6.2 Å (EMD-0366) (Fig. 2A). The binary Impβ:Histone-H1.0 complex was deposited under the model and map accession codes 6N89 and EMD-0367, respectively, with a resolution of 7.5 Å. Both atomic models exhibited truncated side chains (up to CB or CA atoms) and demonstrated poor fit to their respective experimental cryo-EM maps (Fig. 2C, Supplementary Fig. S5, Supplementary Table S6). To address these limitations, these two structural models (Imp7:Impβ:Histone-H1.0 and Impβ:Histone-H1.0) were extensively re-refined through alternating cycles of Rosetta model-map fitting tools ^6^ and Phenix.real_space_refine ^7^ to reassess their alignment with the experimental cryo-EM map. It shall be noted that in the initial refinement step, Rosetta reconstructed all missing side-chain atoms generating a full-atom model that was subsequently fitted to the corresponding cryo-EM map. The same extensive refinement procedure, combined with additional manual model adjustments, was applied to AlphaFold3 predicted complexes with swapped positions of Imp7 and Impβ: Impβ:Imp7:Histone-H1.0^AF3^ and Imp7:Histone-H1.0^AF3^ yielding revised structural models of Impβ:Imp7:Histone-H1.0 (PDB_id 9QEJ) and Imp7:Histone-H1.0 (PDB_id 9QF0) complexes. The refinement was facilitated by manual model rebuilding using Coot, with progress monitored through cryo-EM validation tools, including Map-Model FSC curves, FSCaverage, Map-Model Correlation Coefficient and TEMPy and as implemented in the CCP-EM suite ^8–10^ and Phenix (Supplementary Table S6). This iterative refinement approach enabled a more precise evaluation of the model fit, revealing that the positions of Imp7 and Impβ had been swapped in the original cryo-EM models (6N88 and 6N89) (Fig. 1, 2, Supplementary Fig. S5). This likely arose from the limited resolution of the cryo-EM maps and the lack of comprehensive model validation against available cross-linking ^11^ and biochemical data. Crucially, the cross-linking data unequivocally demonstrate that the H1 core domain, particularly the WGH motif, interacts exclusively with Imp7 (Fig. 1B, C) and not with Impβ. Docking of full-length Impβ crystal structures into the cryo-EM map (EMD-0366) using the phenix.dock_in_map algorithm yielded a striking finding (Supplementary Fig. S6). Four distinct Impβ models (PDB IDs_chain-ID: 1ukl_A, 1ukl_B, 2q5d_B, 8gcn_A) were unequivocally positioned within the region previously assigned to Imp7. In contrast, two Impβ structures: 2qna_A (bound to the IBB domain of Snurportin1) and 3w5k_A (bound to the C2H2 zinc-finger protein SNAIL1), aligned with the contradictory assignment, underscoring the necessity of rigorous structural validation. Remarkably, the earlier assignment was based on the latter Impβ structure (3w5k_A), which, due to its interaction with SNAIL1 (a molecule of similar length to the histone), adopted a conformation deviating from its characteristic curved solenoid architecture observed in other Impβ structures.

---

**Figure S1:**
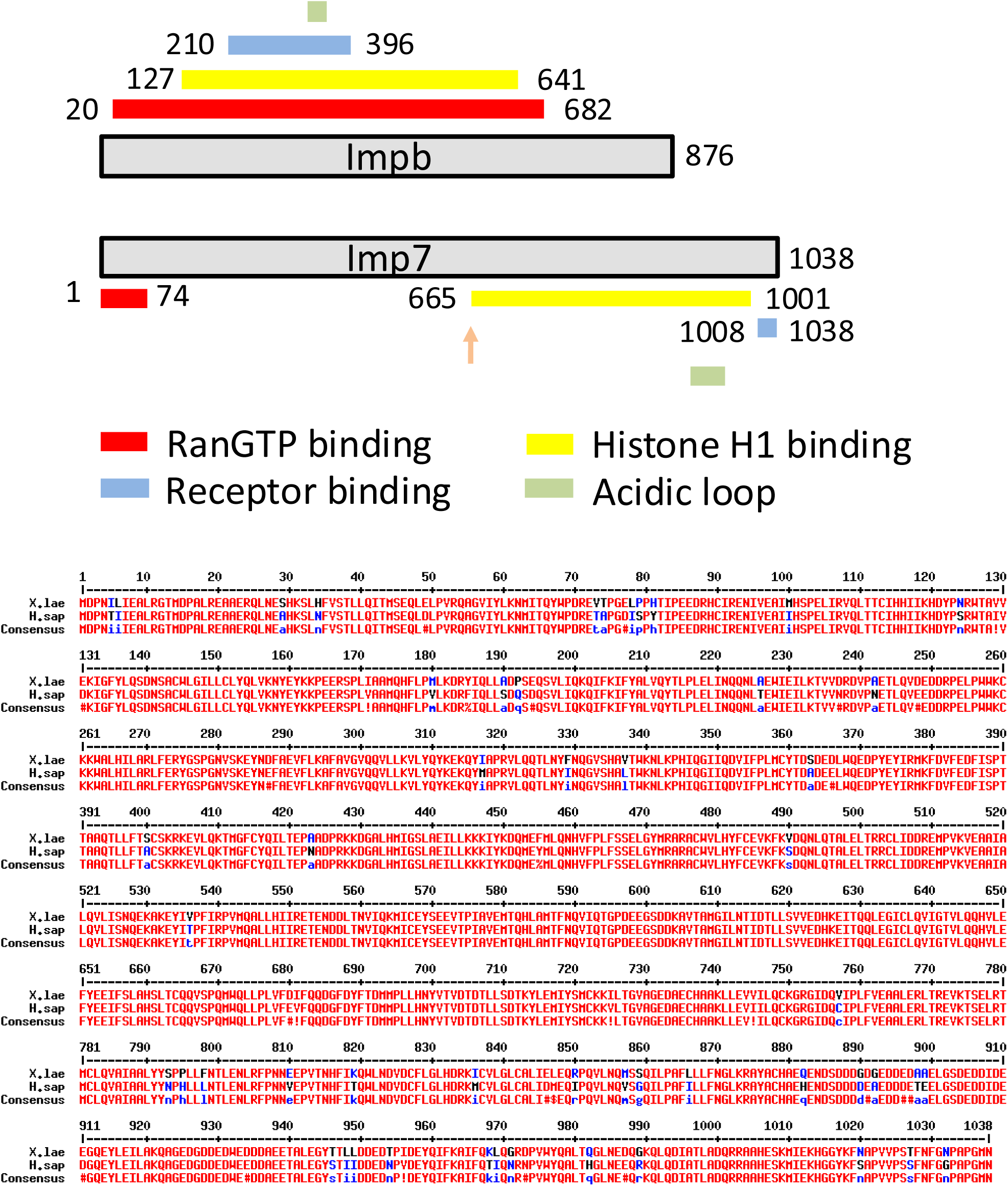
An overview of the structural organization of Impβ and Imp7. In the upper part, the linear arrangement of binding domains in both transport receptors is depicted, with the specific residues marking the beginning and end of each domain noted. This illustrates the domain architecture relevant to their function and interactions. **The lower part** presents a sequence alignment of Imp7 from *Homo sapiens* and *Xenopus laevis*. The alignment highlights the high degree of conservation between the two species, revealing only a few point mutations. This conservation suggests a strong evolutionary pressure to maintain the functional integrity of Imp7 across species.

**Figure S2:**
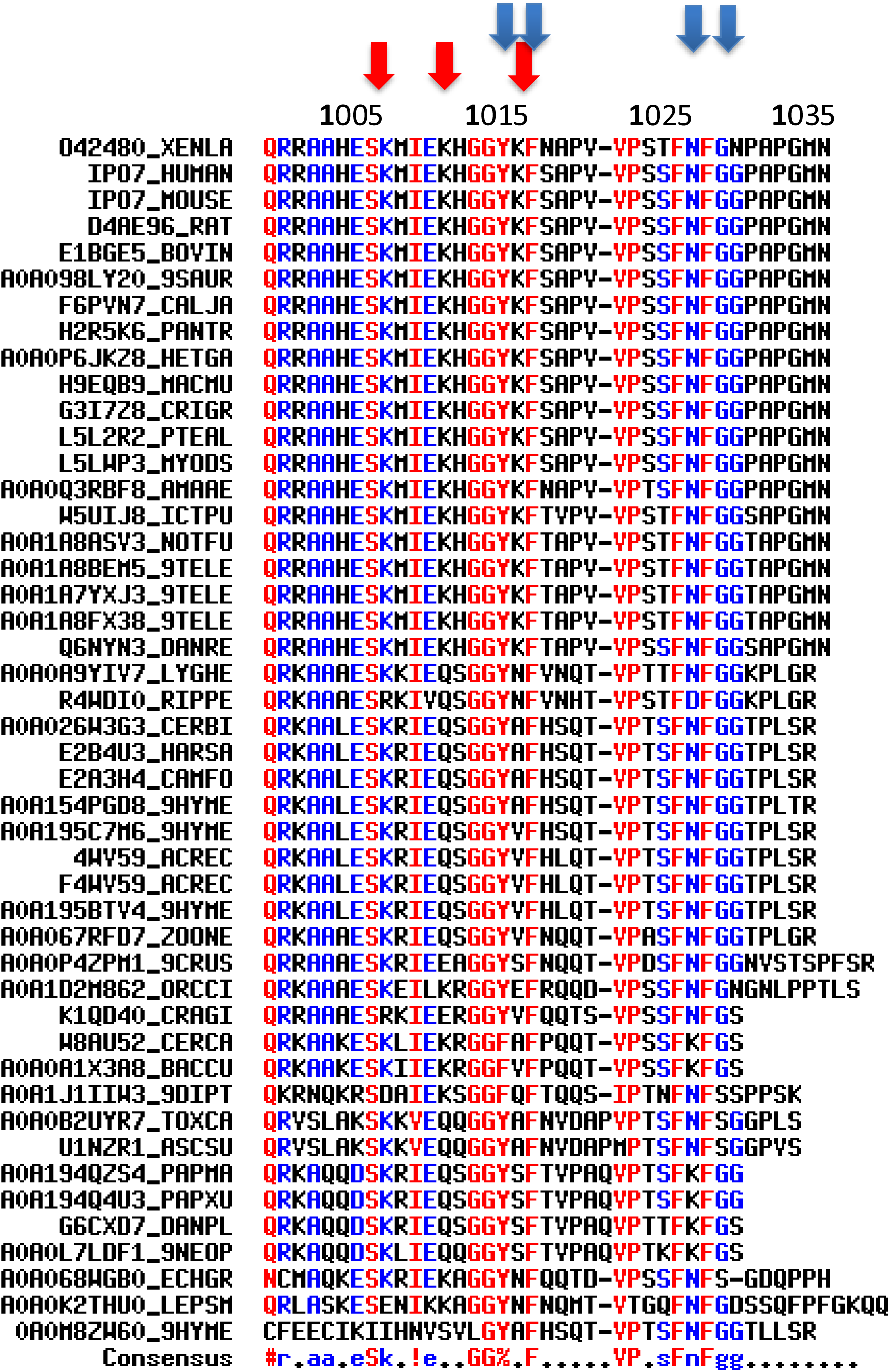
Conservation of NlB (Nucleoporin-like binding) region and indication of the chosen point mutations. Identical residues are marked in red, while highly conserved residues are shown in blue. Conserved lysines presumed essential for function are indicated with red arrows, whereas highly conserved aromatic residues required for binding are marked with blue arrows. Below the alignment, the UniProt references for the Importin 7 sequences used in the alignment are listed. The alignment was performed using MultAlin with standard settings. (Multiple sequence alignment with hierarchical clustering; F. CORPET, 1988, Nucl. Acids Res., 16 (22), 10881-10890). O42480_XENLA: Xenopus lavis (African clawed frog); IPO7_HUMAN: Homo sapiens; IPO7_MOUSE: Mus musculus; D4AE96_RAT: Rattus norvegicus (norvegian rat); E1BGE5_BOVIN: Bos Taurus (cattle); A0A1J1IIW3_9DIPT: Clunio marinus (marine midge); F6PVN7_CALJA: Callithrix jacchus (White-tufted-ear marmoset); K1QD40_CRAGI: Crassostrea gigas (Pacific oyster); A0A0A9YIV7_LYGHE: Lygus hesperus (Western plant bug); H2R5K6_PANTR: Pan troglodytes (Chimpanzee); W8AU52_CERCA: Ceratitis capitata (Mediterranean fruit fly); A0A0B2UYR7_TOXCA: Toxocara canis (Canine roundworm); A0A0A1X3A8_BACCU: Bactrocera cucurbitae (Melon fruit fly); A0A0P4ZPM1_9CRUS: Daphnia magna (Big water flea); A0A026W3G3_CERBI: Cerapachys biroi (Ant); E2B4U3_HARSA: Harpegnathos saltator (Jerdon’s jumping ant); A0A0P6JKZ8_HETGA: Heterocephalus glaber (Naked mole rat); W5UIJ8_ICTPU: Ictalurus punctatus (Channel catfish); H9EQB9_MACMU: Macaca mulatta (Rhesus macaque); Q6NYN3_DANRE: Danio rerio (Zebrafish); A0A068WGB0_ECHGR: Echinococcus granulosus (Hydatid tapeworm); R4WDI0_RIPPE: Riptortus pedestris (Bean bug); 4WV59_ACREC: Acromyrmex echinatior (Panamanian leafcutter ant); A0A194Q4U3_PAPXU: Papilio xuthus (Asian swallowtail butterfly); G6CXD7_DANPL:Danaus plexippus (Monarch butterfly); G3I7Z8_CRIGR: Cricetulus griseus (Chinese hamster); R4WDI0_RIPPE: Riptortus pedestris (Bean bug); A0A194QZS4_PAPMA: Papilio machaon (Old World swallowtail); F4WV59_ACREC: Acromyrmex echinatior (Panamanian leafcutter ant); A0A067RFD7_ZOONE: Zootermopsis nevadensis (Dampwood termite); L5L2R2_PTEAL: Pteropus alecto (Black flying fox); E2A3H4_CAMFO: Camponotus floridanus (Florida carpenter ant); A0A154PGD8_9HYME: Dufourea novaeangliae (sweet bee); L5LWP3_MYODS: Myotis davidii (David’s myotis); A0A0Q3RBF8_AMAAE; Amazona aestiva (Blue-fronted Amazon parrot); A0A195BTV4_9HYME: Atta colombica (leafcutter ant); A0A0L7LDF1_9NEOP: Operophtera brumata (winter moth); A0A195C7M6_9HYME: Cyphomyrmex costatus (Fungus-growing ant); A0A1D2M862_ORCCI: Orchesella cincta (Springtail); U1NZR1_ASCSU: Ascaris suum (Pig roundworm); A0A098LY20_9SAUR: Hypsiglena sp. JMG-2014 (Nightsnake); A0A1A8ASV3_NOTFU: Nothobranchius furzeri (Turquoise killifish); A0A0K2THU0_LEPSM: Lepeophtheirus salmonis (Salmon louse); A0A1A8BEM5_9TELE: Nothobranchius kadleci (Kadlec’s killifish); A0A1A7YXJ3_9TELE: Aphyosemion striatum (Red-striped Killifish); A0A1A8FX38_9TELE: Nothobranchius korthausae (Kinungamkele red tail killifish).

**Figure S3:**
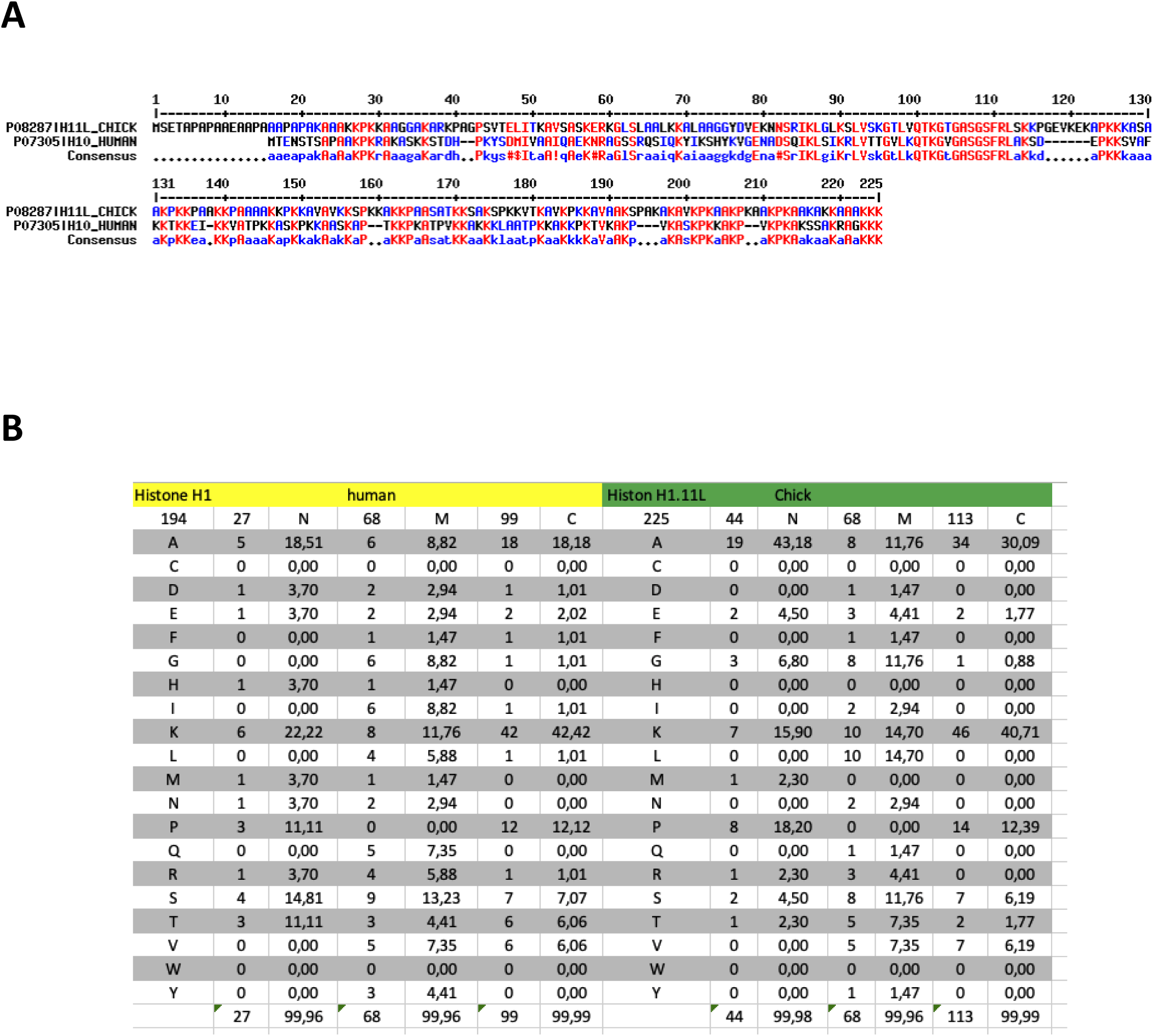
Alignment H1 from *Gallus gallus* and *Homo sapiens*. **A.** The sequences of ggH1.11L (UniProt id: P08287) and hsH1.0 (UniProt id: P07305) were aligned using Multalin (Multiple sequence alignment with hierarchical clustering; F. CORPET, 1988, Nucl. Acids Res., 16 (22), 10881-10890). **B.** Comparison of amino acid composition of the two proteins by region (N-terminal flexible region: N, core domain: M, C-terminal tail: C)

**Figure S4:**
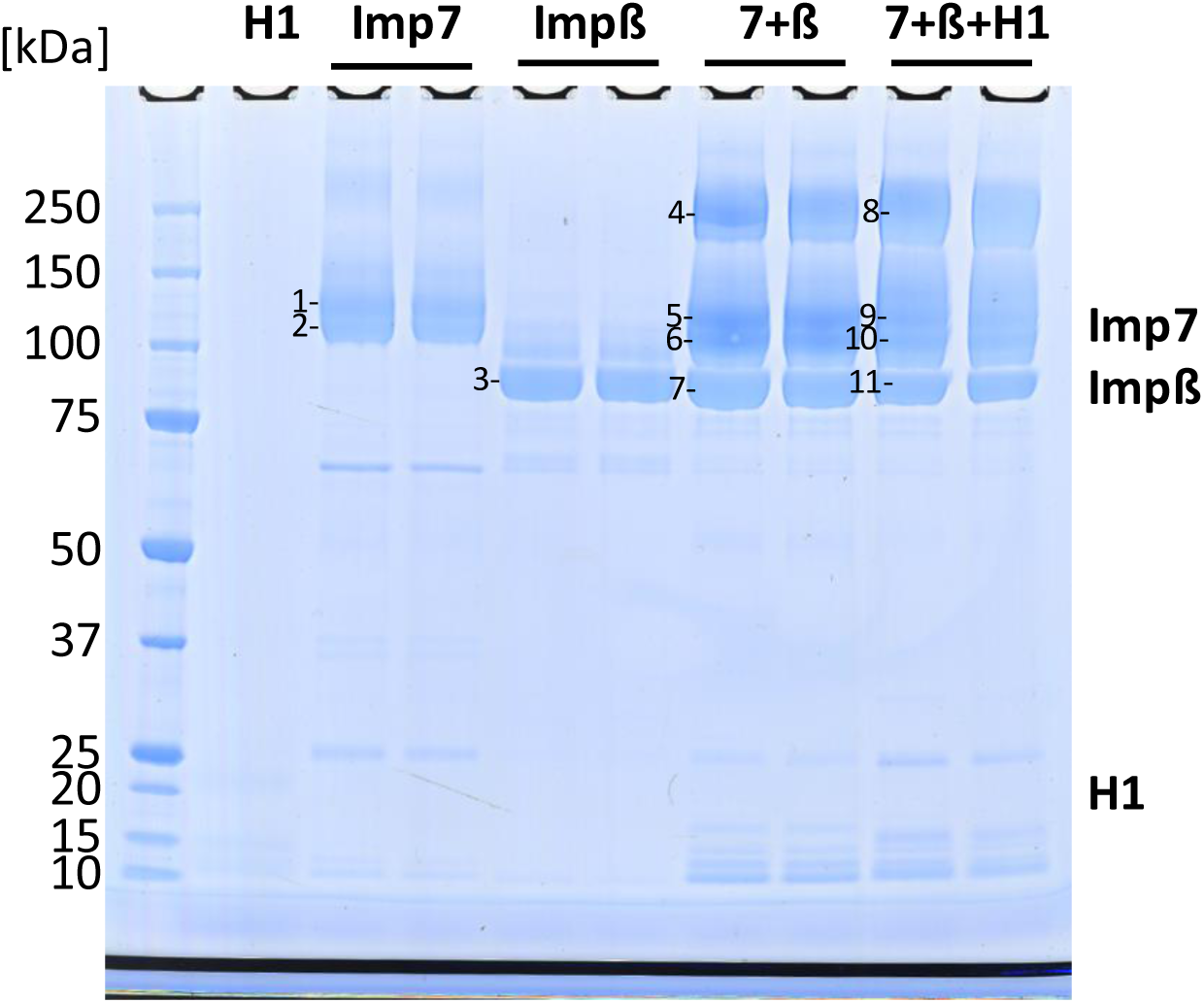
SDS-polyacrylamide gel of BS3-cross-linked histone 1.11L, importin 7 and importin ß and complexes thereof. xlImp7, hsImpβ and ggH1.11L, purified as described in Wohlwend et al. 2007, were subjected to protein-protein cross-linking mass spectrometric analysis (CXMS) as described in main text materials and methods section. The BS3-cross-linked proteins and complexes thereof were separated using SDS-PAGE and visualized by Coomassie staining. The numbered bands were excised and analyzed by CXMS. See Supplementary Table S3 for results. *Wohlwend, D., Strasser, A., Dickmanns, A., Doenecke, D. & Ficner, R. Thermodynamic analysis of H1 nuclear import: Receptor tuning of importinβ/importin7. *Journal of Biological Chemistry 282, 10707–10719* (2007)

**Figure S5:**
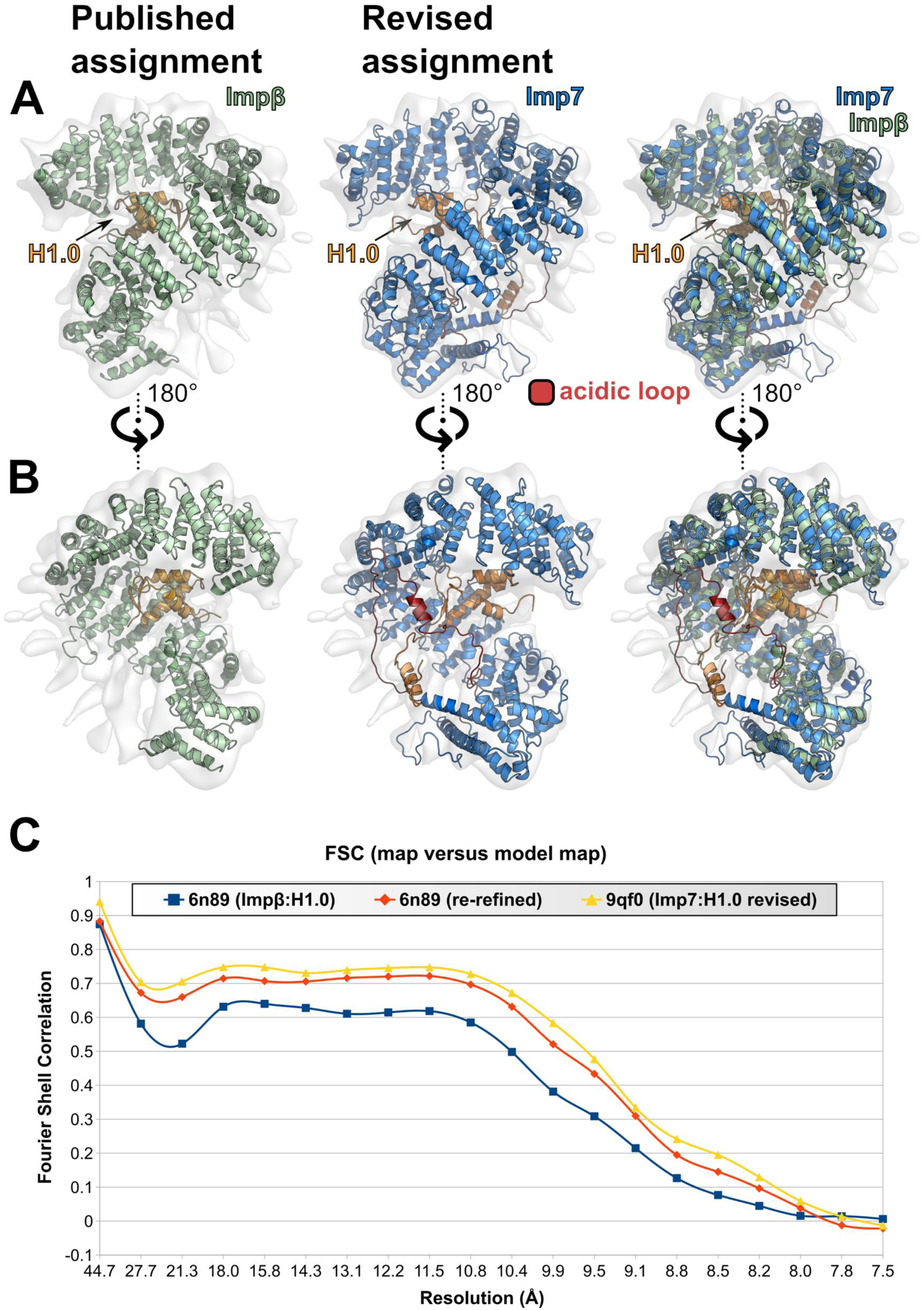
Comparison of the newly fit Imp7:H1 complex and comparison to the original Impβ:H1 structure reveals its better fit. **A.** Panels from left to right: Impb:H1 (PDBid: 6N89), Imp7:H1 and overlay of both. In each case the map (EMDBid: 0367) is shown in light gray. **B.** 180° turn of the orientation in A. **C.** Fourier shell correlation versus resolution of the structures of Impβ:H1, a re-refined version of Impβ:H1 (as described in main text materials and methods section) and the Imp7:H1 based on the AF3 model.

**Figure S6:**
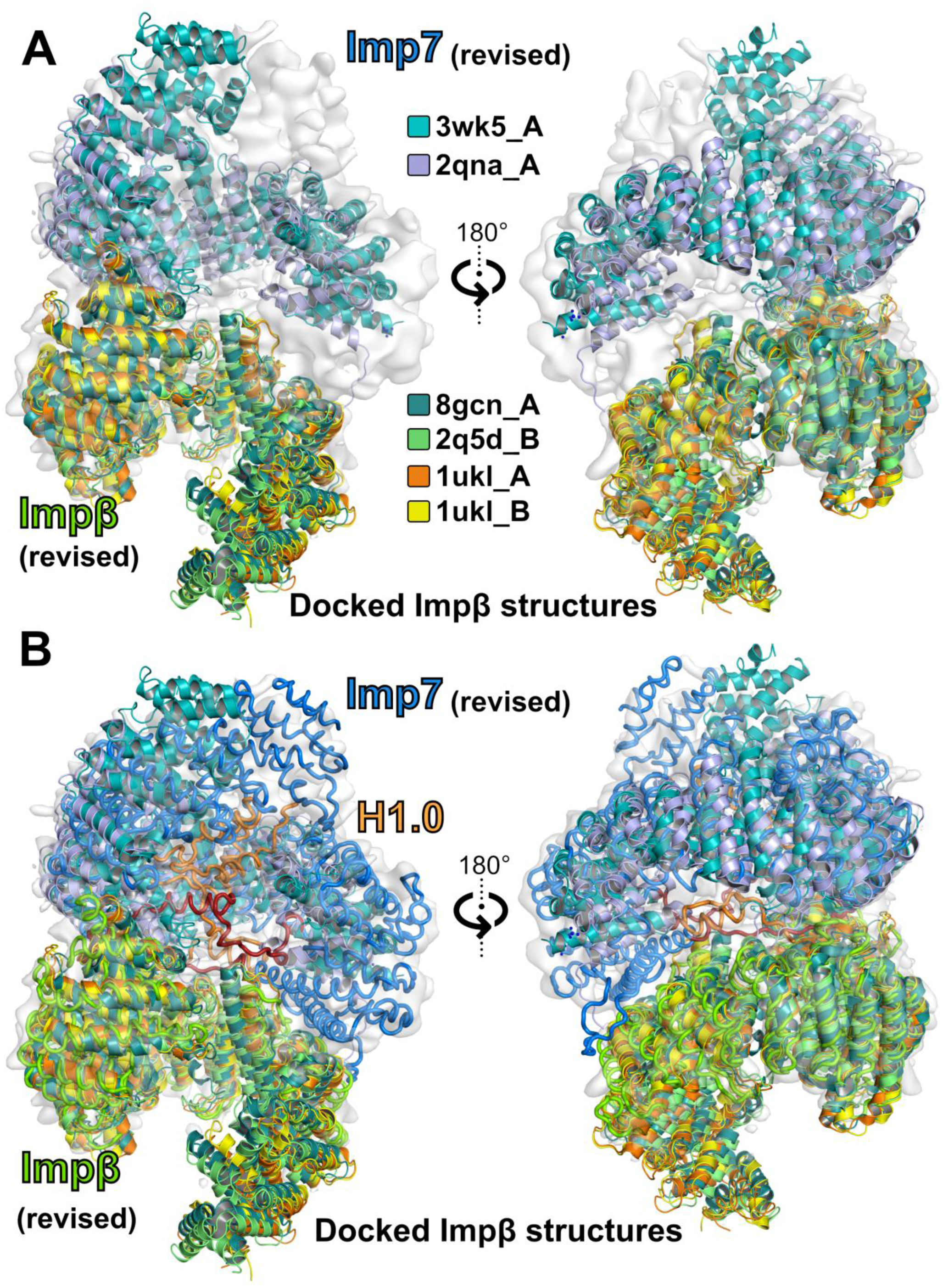
Deposited crystal structures of Impβ were used for automated placement into the EM-map. The phenix.dock_in_map program to dock full-length Impβ crystal structures into the cryo-EM map (EMD-0366) revealed a striking observation: four models (PDB IDs_chain-id: 1ukl_A, 1ukl_B, 2q5d_B, 8gcn_A) were unequivocally placed in the region previously misassigned as Imp7. In contrast, two Impβ structures: 2qna_A (bound to the IBB domain of Snurportin1) and 3w5k_A (bound to the C2H2 zinc-finger protein SNAIL1), aligned with the incorrect assignment, highlighting the importance of rigorous structural validation.

**Figure S7:**
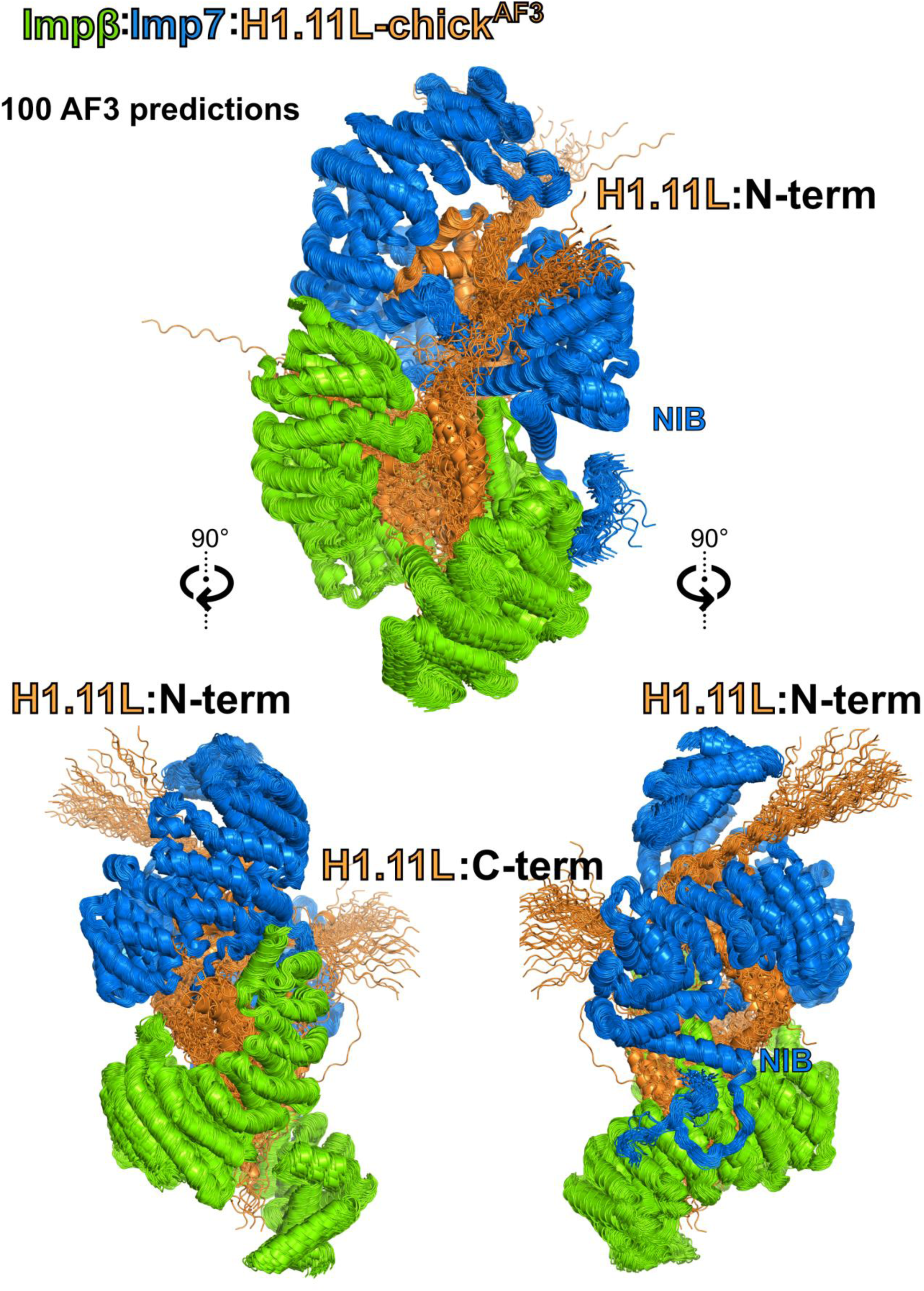
AF3-modelling fitted the H1 core into Imp7 and the H1-Lys rich flexible regions within **Impβ.** Interestingly, the termini predominantly exit the central cavity by interactions on Imp7.

**Figure S8:**
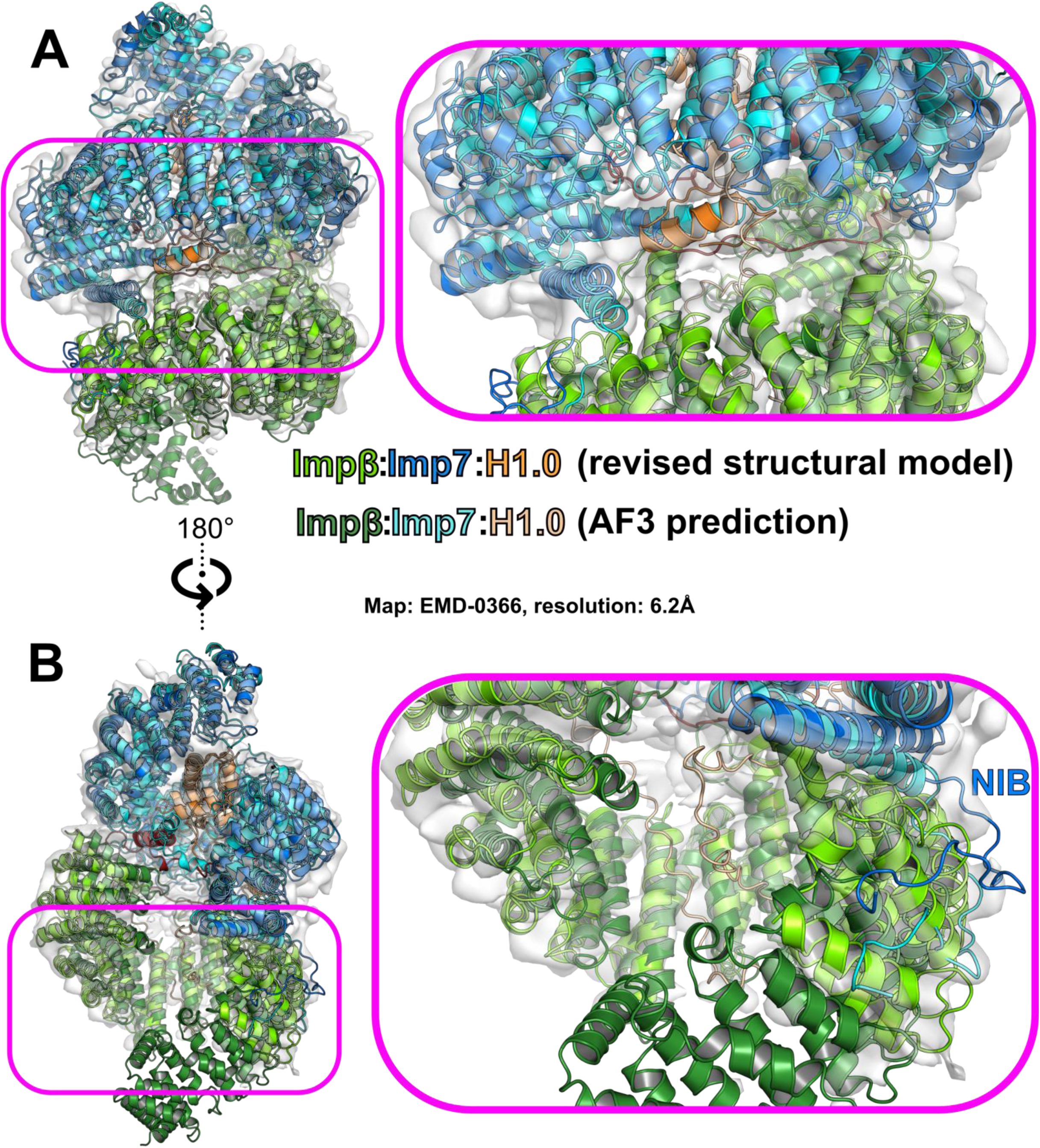
Comparison of the AF3 generated model of Impβ:Imp7:H1 and the refined structural model. **A.** Overall structure of the ternary complex (left) and magnification of the boxed region (right) reveals only subtle changes at the interface of the three proteins. In contrast, in **B.** is presented the 180° rotational view (right) and the magnification of the NlB region and its binding site, which are more compact in the structural model. Coloring of the individual proteins as indicated in the figure.

**Figure S9:**
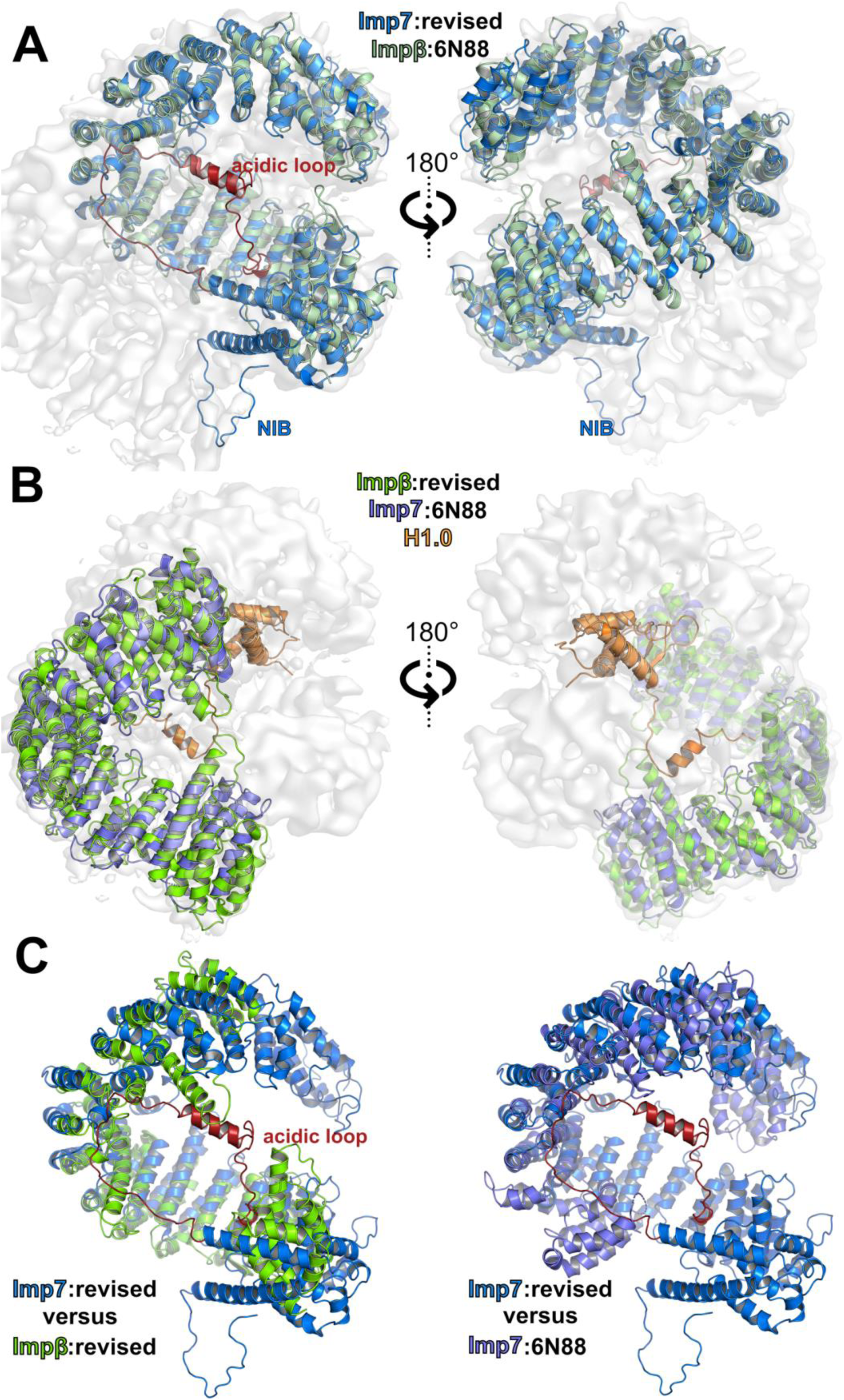
Comparison of the Imp7:Impβ:H1 complex (6N88) and the revised structural model Impβ:Imp7:H1 against the cryo-EM map (EMDB-0366). **A.** Superposition of the respective molecules positioned in the upper part of the map in two orientations. **B.** Superposition of the respective molecules positioned in the lower part of the map in two orientations. **C.** The overlay of Imp7 and Impβ of the revised structure (left panel) as well as the overlay of Imp7 of the revised structure and Imp7 from 6N88 (right panel) exhibit a similar curvature and arrangement in the middle region, but distinct differences C– and N-terminal regions.

**Figure S10:**
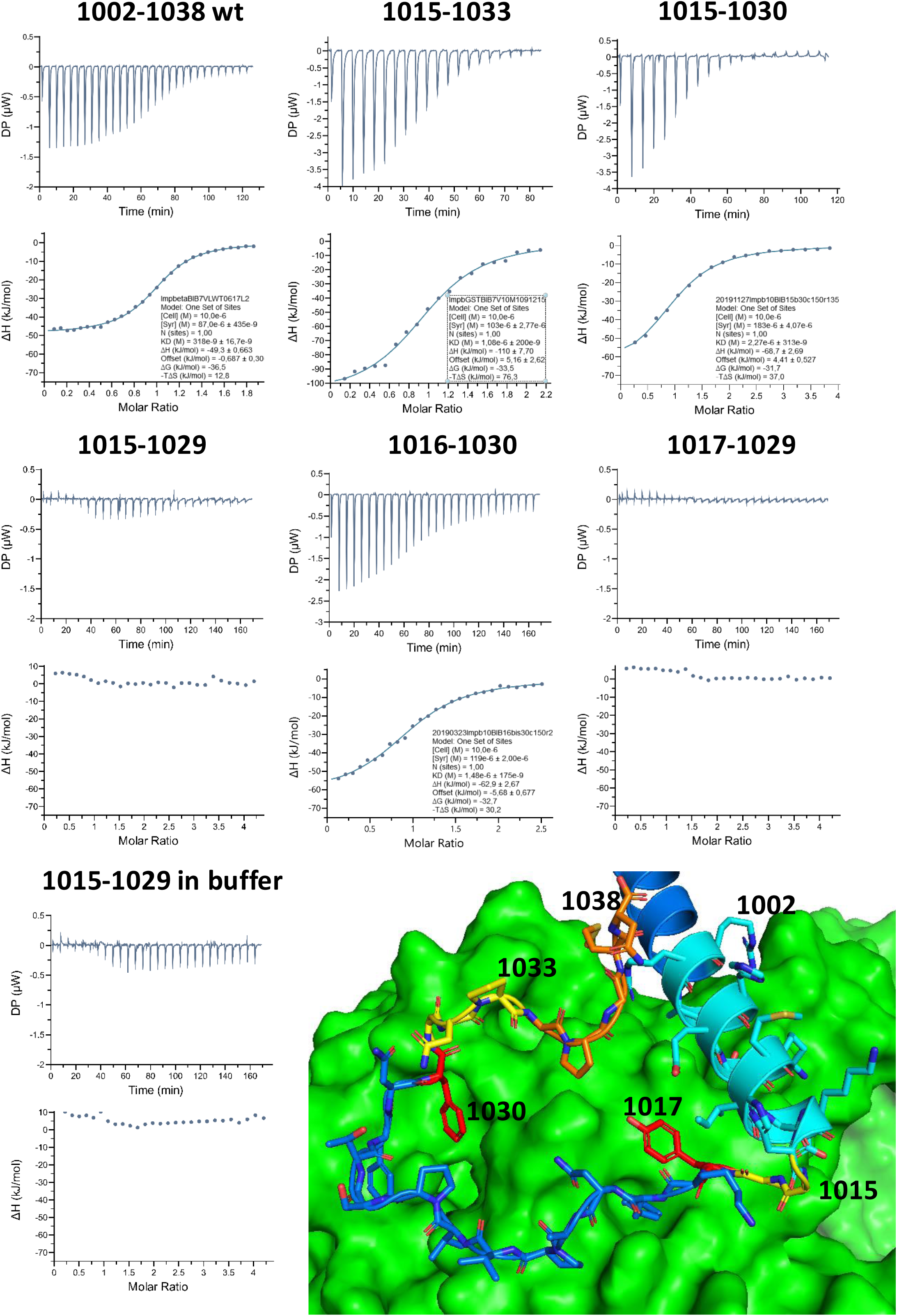
ITC results of the tested deletion mutants reveal the requirement of residues 1016-1030 for binding to Impβ. The thermograms and data evaluation of representative experiments, selected to be close to the average values reported in Table 1 of the main text, are shown. The results for the deletion constructs are shown, with the numbers above indicating the regions expressed as GST-fusion proteins. These fusion proteins contain a PreScission Protease cleavage site and an additional GS dipeptide due to the restriction site used for cloning, positioned before the actual first residue of the NlB-region in order to improve accessibility. The bottom right panel depicts the complete NlB-region starting at residue 1002 (cyan). Yellow residues highlight the GG-pair marking the beginning of the flexible tail region of the NlB-region, as well as C-terminal residues that could be deleted. The stretch shown in blue, flanked by red aromatic residues, represents the required region for binding to Impβ. Additionally, before unpredicted residues from the preceding α-helix are depicted in blue.

**Figure S11.**
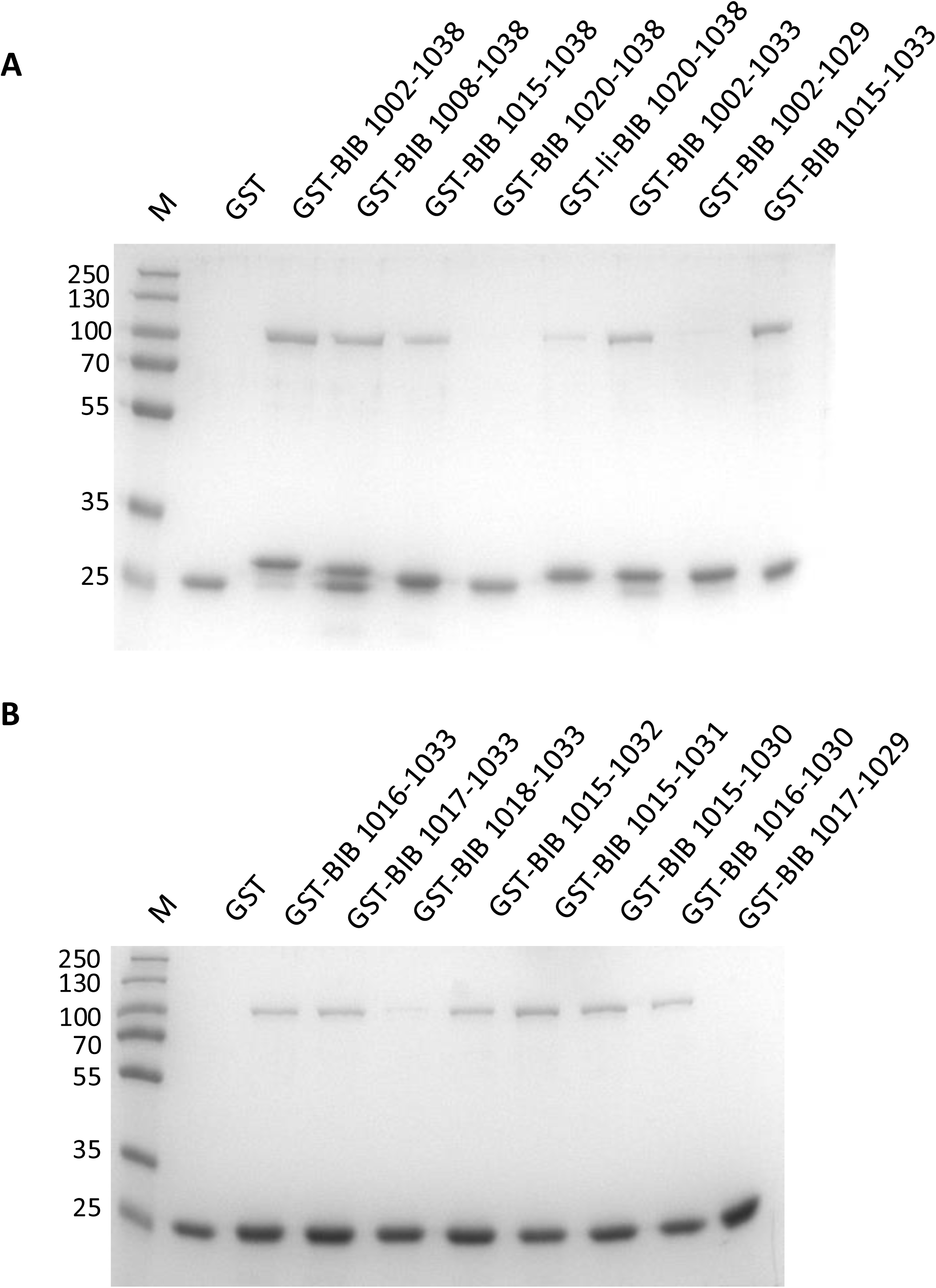
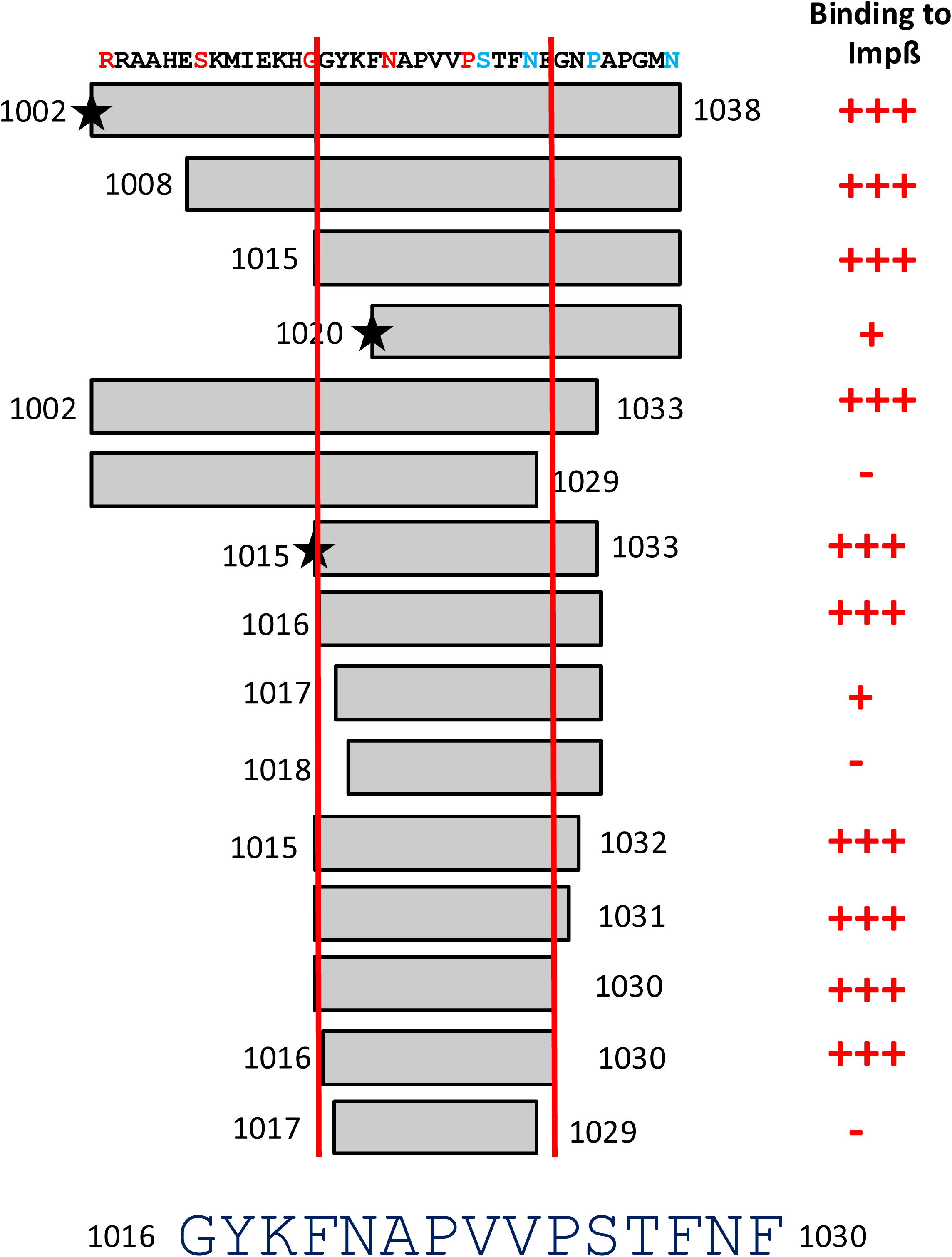
pull down experiments confirm the results obtained from deletions tested in the ITC experiments. **A. & B.** Pulldown experiments were performed as described, analyzed by SDS-PAGE chromatography and visualized by Coomassie staining. **C.** Qualitative summary of the deletion experiments. Detailed description: Deletion mutants of the NlB-domain were constructed resulting in GST-fused fragments, expressed purified (M&M) and used in pull-down experiments with Impβ (for a comprehensive list see Table. S1). Deletion of the first six residues (NlB1008-1038; the deletion mutant form used in (Bäuerle, 2002) did not alter the binding properties, neither did the deletion of the next seven residues from the N-terminus (NlB1015 – 1038). In contrast, further deletion by the next five or ten residues abrogated binding of Impβ. From the C-terminal end, only the deletion of the last five residues (NlB1002-1033) did not compromise Impβ binding, whereas further deletions (NlB1002-1029, NlB1002-1026) abolished binding completely. A fragment with residue 1033 at the C-terminus and different deletions from the N-terminal side resulted in the expected fragment comprising residues 1015 to 1033 (NlB1015-1033), which is still able to interact with Impβ in pull down experiments. Further deletions revealed that 1016-1030 still binds to Impβ, whereas shorter versions (NlB1020-1033, NlB1025-1033) did not bind anymore. In order to exclude spatial restraints between the GST-moiety and the NlB-fragment as reason for a lack of binding, a 15-residue linker (see M&M in main text) was introduced in some fragments to extend the distance between GST and the NlB fragment. The resulting proteins were also tested and exhibited the same properties as the ones without the linker (Fig. 11C indicated by a star), excluding the possibility of spatial restraints as reason for non-binding to Impβ. In a second round, deletions fragments of Imp7 were cloned, expressed and used in pull-down experiments, that were shortened by steps of one residue either from the N or C-terminal side starting at 1015 and 1033, respectively. Deletion of residues 1015 from the N-terminus or three residues (down to G1031) from the C-terminus did not alter the binding properties significantly (Fig. 1B). As soon as residues 1016 or 1030 (mutant NlB 1017-1029) were removed the binding dramatically decreased. So, it can be concluded, that the minimal binding region lies within residues 1016 to 1030 of Imp7. Bäuerle, M., Doenecke, D. & Albig, W. The requirement of H1 histones for a heterodimeric nuclear import receptor. *Journal of Biological Chemistry* **277**, 32480–32489 (2002).

**Figure S12:**
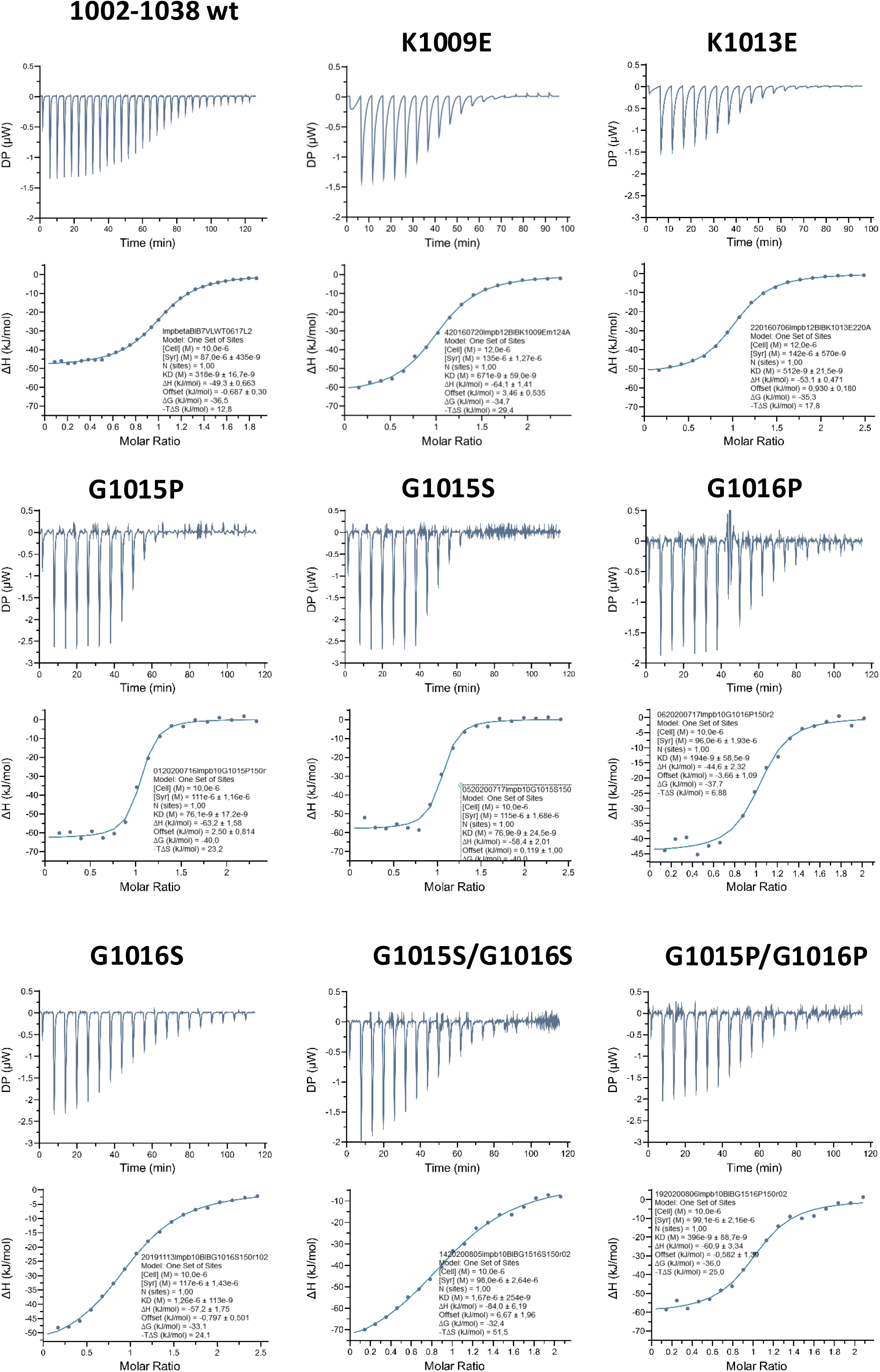
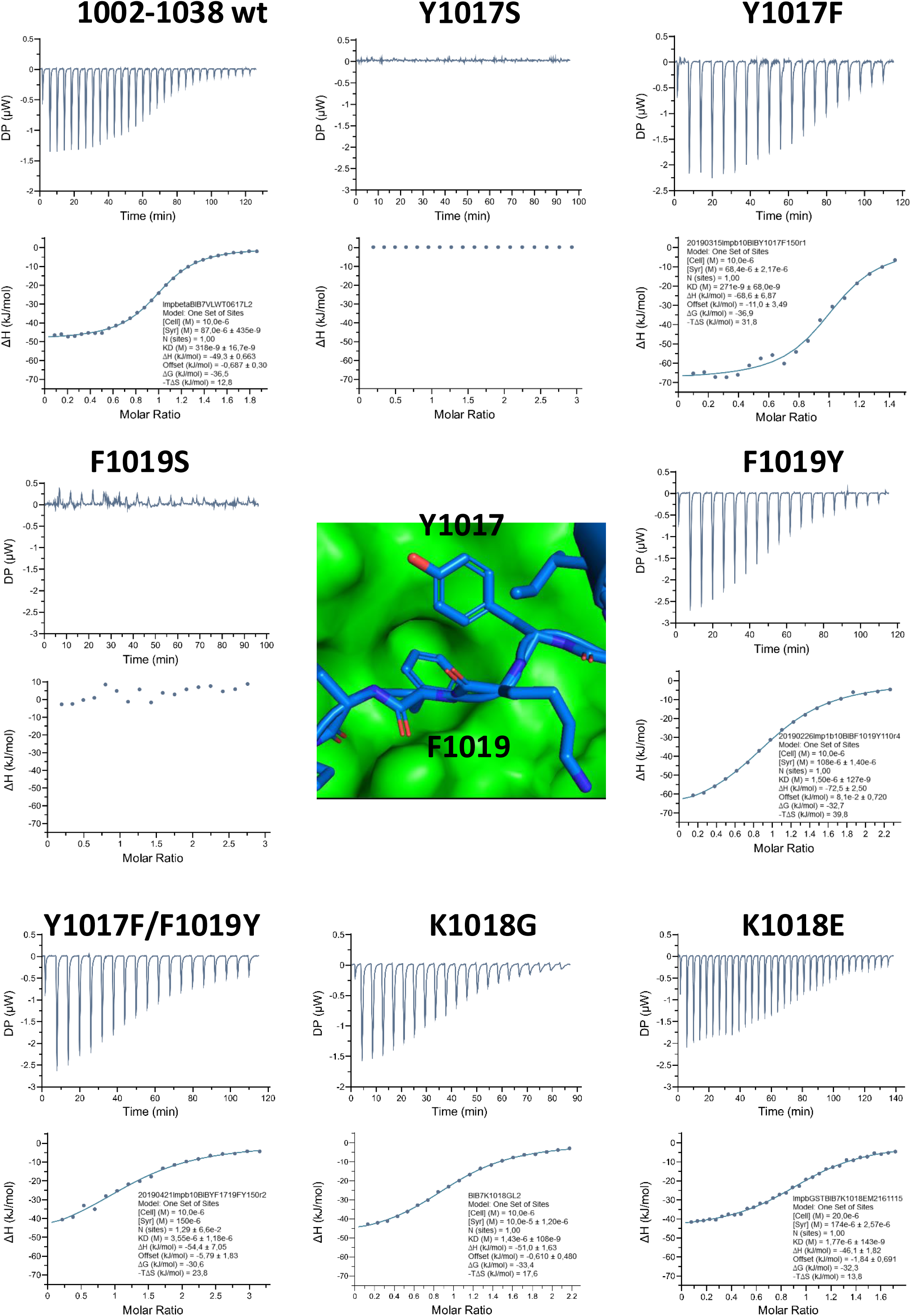
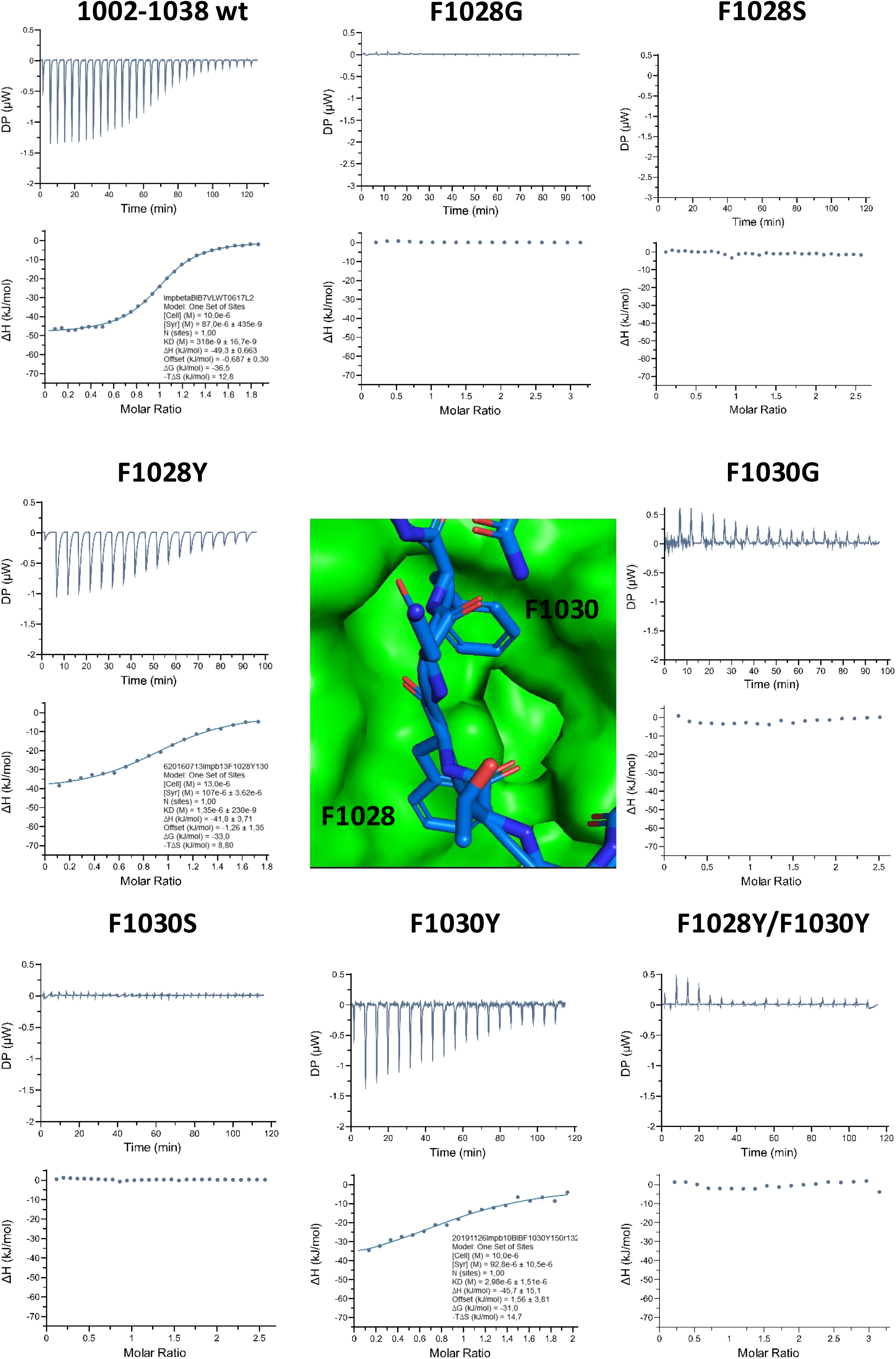
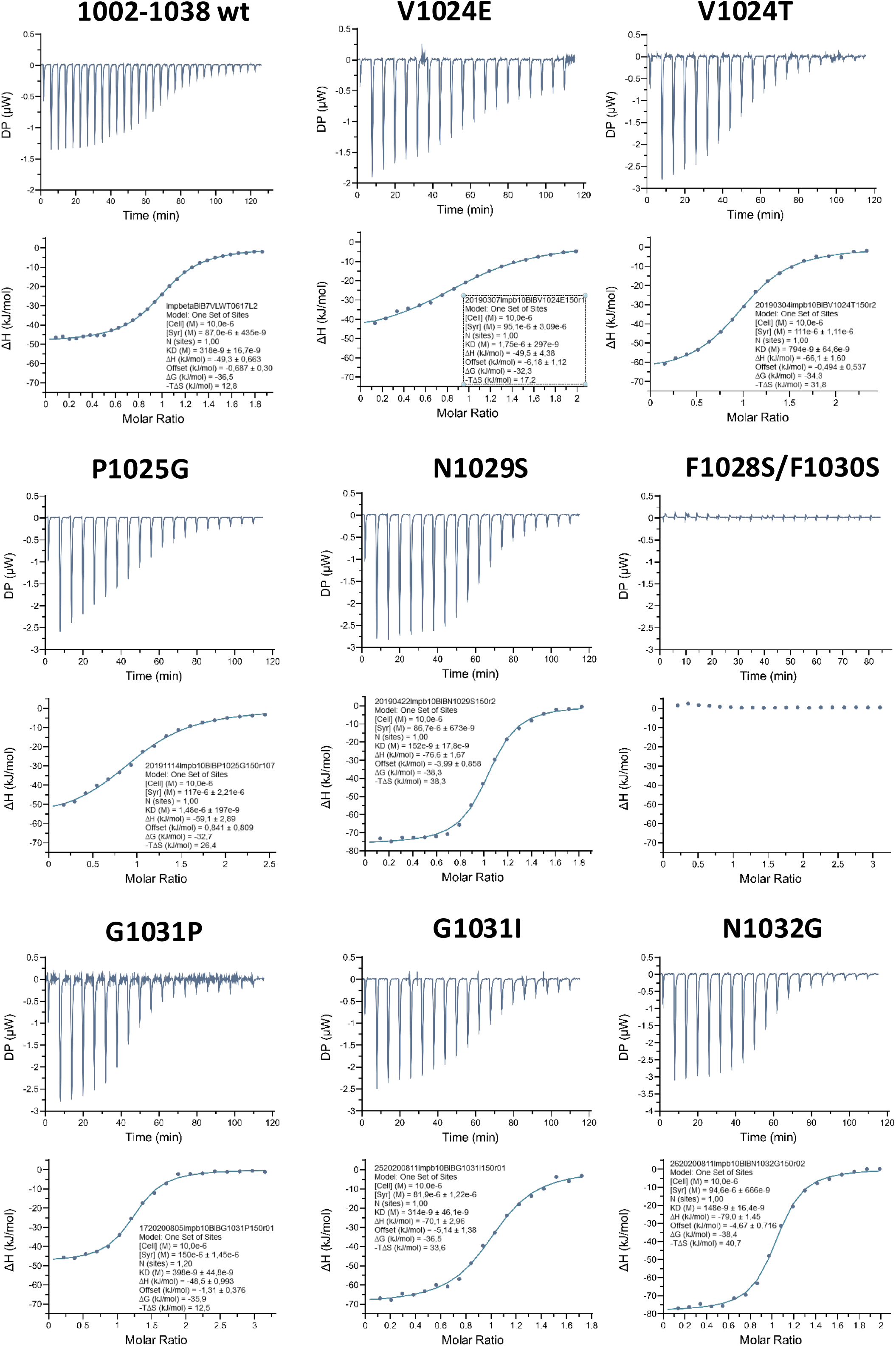
The ITC results of the tested point mutations are presented. The thermograms and data evaluation of representative experiments, selected to be close to the average values reported in Table 2 of the main text, are shown. A. Lysine residues (see also B.) within the NlB region, as well as the GG pair preceding the intrinsically disordered region (IDR), do not play a significant role in binding to Impβ, contrary to early speculations. B. The first binding motif, YxF at positions 1017 and 1019, is required and essential for binding. However, either Y or F at both positions is sufficient to secure binding. The central image illustrates the binding mode (see also main Fig 3). C. The second binding site, FxF, is required and essential for binding. At most, only one F residue may be substituted with Y at a time to retain binding. The central image illustrates the binding mode (see also main Fig 3). D. The remaining conserved residues (Fig S2) affect binding affinity but do not completely abolish the interaction between the NlB region and Impβ.

**Figure S13:**
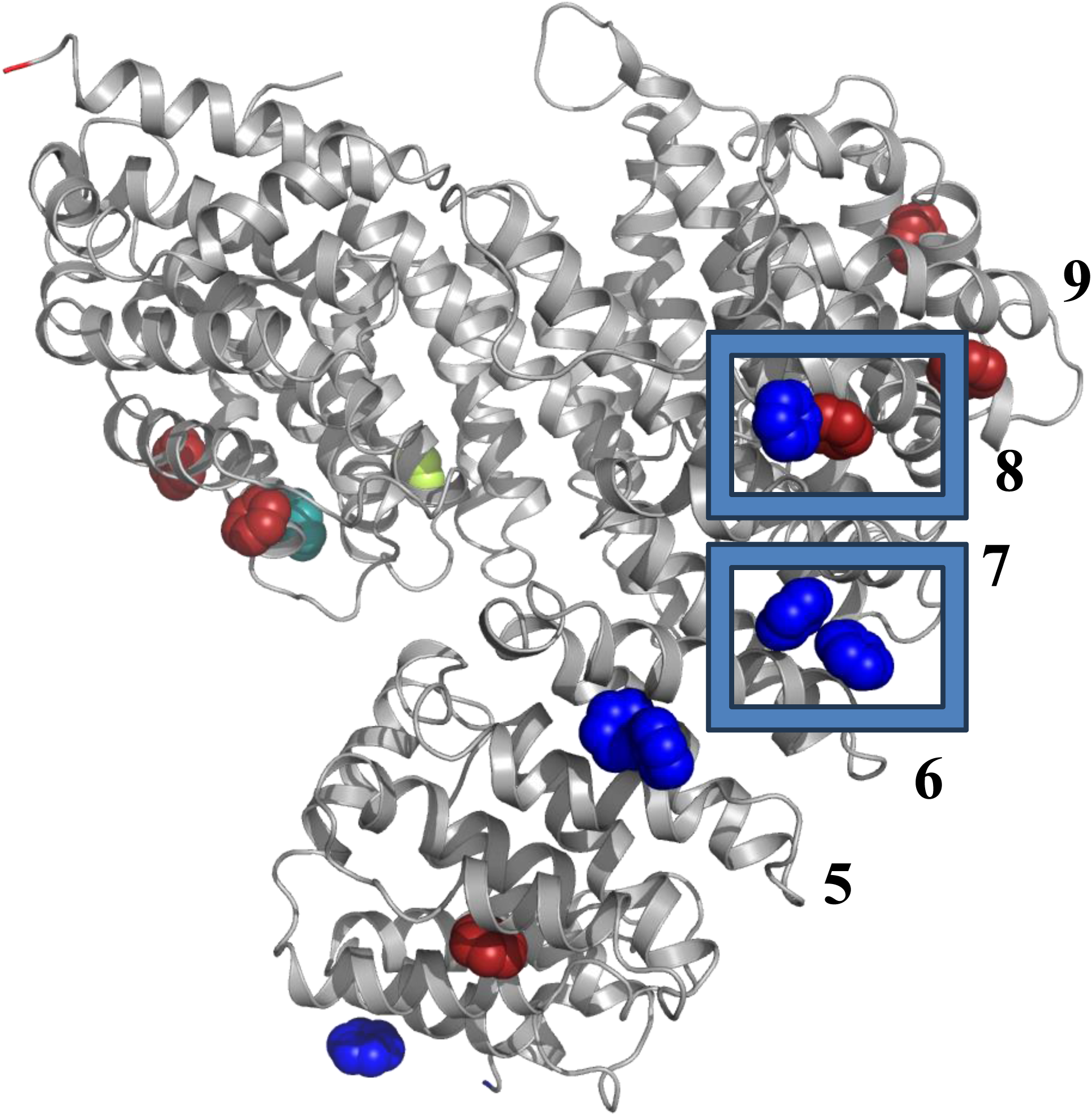
Experimentally (X-Ray and biochemically) defined as well as *in silico* predicted FG binding sites on the surface of Impβ. The coloring indicates the following: Blue: found in crystal structures, Dark red: found by *in silico* simulation, Green: biochemically determined, Limon: both in *silico* and biochemically. The numbering indicates the A helices of the HEAT repeats forming the respective cleft for NlB region binding.

**Figure S14:**
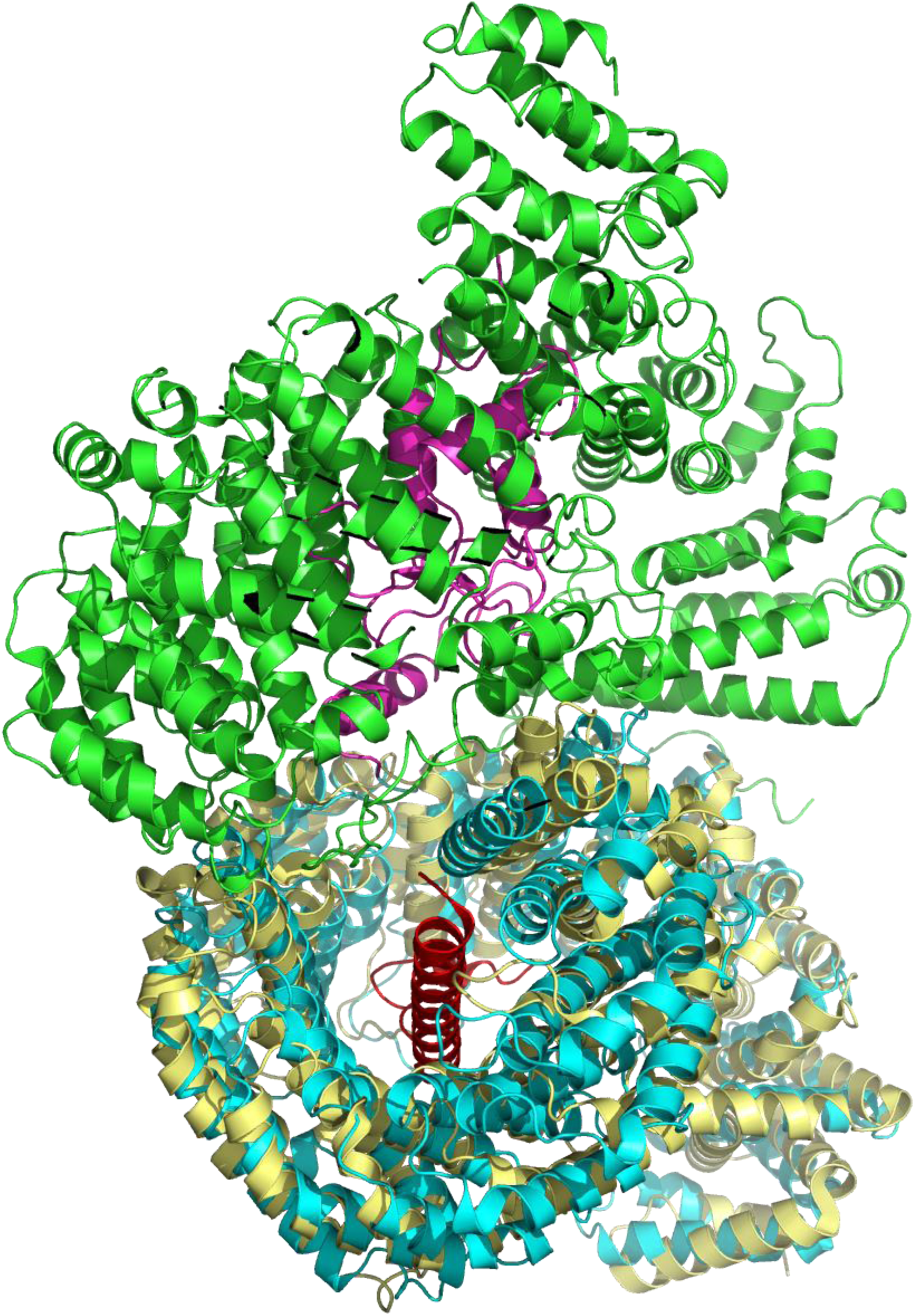
The Impβ:Imp7:H1 complex is still able to interact with the IBB domain of Impα. The structure of Impβ:Impα−IBB (PDBid: 8gcn) was superposed to Impβ of the ternary complex. Binding of H1 and Impα−IBB do not interfere with each other in this arrangement, as shown experimentally by Jäkel et al. (Jäkel, S. et al. The Importin β/Importin 7 Heterodimer Is a Functional Nuclear Import Receptor for Histone H1. The EMBO Journal vol. 18 https://www.embopress.org (1999))

**Figure S15:**
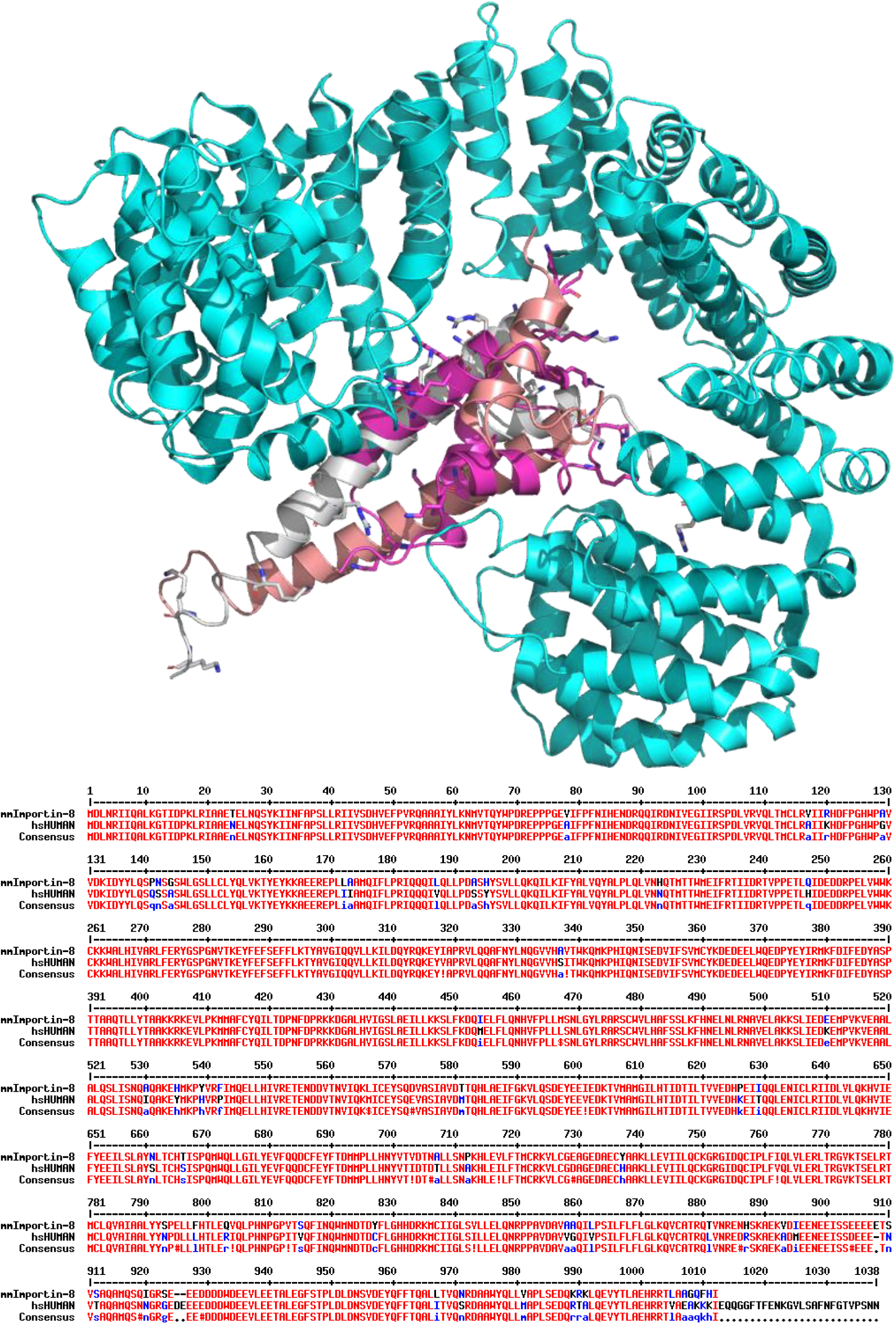
H1 core and SREBP could bind to Impβ similarly. **A.** The structure of Impβ bound to a dimeric fragment of SREBP (PDBid: 1UKL), was used to manually superimpose the hsH1-core region. **B.** Alignment of Impβ: *mus musculus* versus *homo sapiens*. The sequences of hsImpβ (UniProtid: Q14974) and mmImpβ (UniProtid: P70168) were aligned using Multalin revealing very little differences (Multiple sequence alignment with hierarchical clustering; F. CORPET, 1988, Nucl. Acids Res., 16 (22), 10881-10890).

